# USP18 Inhibition Enhances Type I Interferon Signalling and Immune Activation in the Tumour Microenvironment of Triple-Negative Breast Cancer

**DOI:** 10.1101/2025.09.01.672152

**Authors:** Chartinun Chutoe, Emily Rose Webb, Morwenna Theresa Muir, Fraser Laing, Valerie G Brunton, Alexander von Kriegsheim

**Author notes:** To whom correspondence should be addressed: Alexander von Kriegsheim.

## Abstract

Triple-negative breast cancer (TNBC) is one of the most aggressive and treatment-resistant breast cancers. Although immunotherapy has emerged as a promising treatment option, clinical benefit is limited, with only around half of patients responding, even when combined with standard chemotherapeutic agents. This limited efficacy is often attributed to immunologically “cold” tumour microenvironments (TME), which are resistant to current immunotherapies. Addressing this challenge requires approaches that can reprogram “cold” TMEs into “hot” immune-responsive states.

USP18, a negative regulator of type I interferon (IFN) signalling, suppresses immune activation by removing ISG15 from target proteins and disrupting IFNAR–STAT2 interactions. Here, we show that both genetic ablation and catalytic inactivation of USP18 enhance type I IFN signalling in TNBC cells, leading to sustained STAT1/STAT2 phosphorylation. This increased IFN responsiveness promotes antigen presentation via MHC-I upregulation and increases expression of pro-apoptotic ligands such as FAS.

Proteomic profiling and immunophenotyping revealed that USP18 inhibition *in vivo* reduces tumour growth and increases immunogenicity, accompanied by cancer-immune infiltration modulation including CD8⁺ T cells, Th1 cells, NK cells, cDC1, and pro-inflammatory M1-like macrophages. These changes reflect a shift in the TME from an immunosuppressive to an immunostimulatory state, driven by heightened and prolonged type I IFN signalling.

Our findings highlight the therapeutic potential of USP18 inhibition to convert immunologically “cold” tumours into “hot” tumours, by enhancing IFN-driven immune activation and improving the efficacy of immunotherapy in TNBCs.

## Introduction

Breast cancer is the most common solid tumour in women worldwide, with some subtypes being highly aggressive, resulting in metastasis to distant organs such as bone, lung, and liver ^1–4^. Triple-negative breast cancer (TNBC) has been recognised as the most aggressive subtype ^5–7^. Furthermore, TNBC exhibits higher expression of programmed death ligand-1 (PD-L1) than other subtypes ^8–10^. PD-L1 binds to programmed death receptor 1 (PD-1) on T cells, leading to T-cell exhaustion, contributing to an immunosuppressive, “cold” environment, and ^11–15^ negatively affecting the anticancer immune response in TNBC ^8, 11,13^. Additionally, “cold” immunosuppressed tumours exhibit a poor immune response due to low immune cell infiltration, low immunogenicity due to poor antigen presentation, leading to resistance to immunotherapy ^16^. Strategies to convert “cold” into “hot” tumours, such as combining immune checkpoint inhibitors (ICIs) with chemotherapy or targeted therapies, are being explored to enhance treatment efficacy ^17,18^.

Type I interferon (IFN) signalling is crucial for regulating cellular immunity against infections ^19–21^. This response is triggered upon IFN stimulation when type I IFNs bind to type I IFN receptors, including interferon alpha receptor 1 (IFNAR1) and interferon alpha receptor 2 (IFNAR2) ^22^. This interaction leads to the phosphorylation of Janus kinase 1 (JAK1) and tyrosine kinase 2 (TYK2), which promotes the phosphorylation and activation of signal transducer and activator of transcription 1 and 2 (STAT1 and STAT2) ^22^. In conjunction with IFN regulatory factor 9 (IRF9), it forms a complex known as IFN-stimulated gene factor 3 (ISGF3) ^22,23^. This complex is then translocated into the nucleus, where it functions as a transcriptional activator by binding to IFN-stimulating response elements (ISREs) found in a wide range of IFN-stimulated genes (ISGs) ^22–24^. The proteins encoded by ISGs play crucial roles in modulating both innate and adaptive immunity, inhibiting pathogen replication and survival, and regulating key cellular processes such as proliferation, survival, and cell death ^25^. Therefore, enhancing type I IFN response in cancer is one way to promote an immune-responsive microenvironment.

Analogous to most signalling pathways, IFN signalling output is controlled via feedback regulators. A key negative feedback inhibitor of the type I IFN response is ubiquitin-specific protease 18 (USP18) ^26,27^. USP18 is a key player in suppressing the IFN response through a STAT2-dependent mechanism ^28,29^. Consequently, USP18 loss has been shown to induce interferonopathies, which result in chronic inflammation, microcephaly and neonatal death ^30,31^. Point mutations of USP18, substituting isoleucine (I) for asparagine (N) at amino acid position 60, USP18^I60N^, result in a less severe phenotype, triggering IFN-mediated autoinflammation in patients ^32^. The I60N mutation is enzymatically competent but disrupts STAT2-binding^28^, highlighting the importance of the non-enzymatic negative feedback inhibition ^28,32^ in dampening IFN signalling.

Apart from its role in type I IFN suppression, USP18 functions as a deISGylase, a protease that removes the interferon-stimulated gene 15 (ISG15) modifications from proteins ^26,33,34^. ISGylation and deISGylation of proteins have been shown to regulate type I IFN response and breast cancer progression ^35,36^. Substitution of the active-site cysteines to serine residues effectively kills USP18 deISGylase activity, but by itself is not sufficient to trigger IFN sensitivity ^28^. Conversely, substituting the catalytically active site of USP18 with bulkier arginine residues (C64R/C65R) impairs enzymatic activity and enhances interferon sensitivity in chronic myeloid leukaemia cells ^37^. Suggesting that disruption of the active site has the potential to inhibit both the enzyme activity as well as the USP18-STAT2 complex and could sensitise breast cancer to IFN stimulation.

In this study, we hypothesised that USP18 inhibition could enhance IFN-mediated immune responses in TNBC. To investigate this, we employed CRISPR-Cas9 gene editing to generate USP18 knockout and USP18 mutant models (hUSP18^C64R/C65R^ or mUSP18^C61R/C62R^) that contain bulky residues in the active site to mimic the disruption that a small chemical inhibitor binder could trigger. We then conducted high-throughput proteomic analysis to characterise the changes associated with USP18 inhibition. Additionally, we performed immunophenotyping to assess alterations in the tumour immune microenvironment. Together, these approaches enabled a comprehensive evaluation of the functional impact of USP18 inhibition in TNBC and its potential as a therapeutic target.

## Materials and Methods

### Cell culture and reagents

Human triple-negative breast cancer cells (MDA-MB-231; RRID: CVCL_0062) were cultured in high-glucose Dulbecco’s modified Eagle’s medium (DMEM) (D6429, Sigma-Aldrich, MO, USA) supplemented by 10 % foetal bovine serum (16000044, Gibco, Texas, USA). The mouse triple-negative breast cancer cells (4T1; RRID: CVCL_0125) were maintained in RPMI1640 (11530586, Fisher Scientific, NH, USA) supplemented by 10 % foetal bovine serum (Gibco) and 1% sodium pyruvate (11360070, Gibco). For human natural killer cells (NK-92; RRID: CVCL_2142, CRL-2407, ATCC), they were maintained in MyeloCult™ H5100 (05150, STEMCELL Technologies) supplemented by 12.5% horse serum (16050122, Thermo Scientific, MA, USA), 1% Glutamax (35050061, Thermo Scientific) and 10 *µ*g/L IL-2 (130097742, Miltenyi Biotec, Germany). The cells were sub-cultured according to the manufacturer’s instruction and kept at 37°C with 5% CO2.

To study IFN response, human cell lines were treated with human recombinant interferon alpha, IFN-α2b, (11105-1, pbl assay science, USA). While mouse cell line was treated with mouse recombinant interferon alpha 2, IFN-α2 (14-8312-80, Invitrogen).

### USP18 gene modification

To generate USP18 knock out (USP18^KO^), 5×10^5^ cells were electroporated using SE Cell Line 4D-Nucleofector® X Kit S (V4XC-1032, Lonza). crRNA was annealed with tracrRNA (1072533, IDT) at 95°C before making ribonucleoprotein complex with Cas9 protein (1081059, IDT) at room temperature. crRNAs used in this study were showed in Table S1. The cell pools were then check their USP18 expression by WB and proteomic studies.

For catalytically mutated knock in (USP18^C64R/C64R^/ USP18^C61R/C62R^), 5×10^5^ cells were electroporated using the same conditions as the knockout protocol, with the SE Cell Line 4D-Nucleofector® X Kit S (Lonza). To generate ribonucleoprotein complexes, crRNAs (IDT) were annealed with tracrRNA (IDT) and subsequently complexed with Cas9 (IDT). The HDR template was then added to the electroporation mixture. Following electroporation, cells were cultured at 32°C in medium supplemented with HDR enhancer (IDT, 10007921). After 24 hours, the medium was replaced with fresh medium, and cells were maintained at 32°C for an additional 24 hours before being shifted to 37°C. Single-cell clones were selected based on DNA sequencing results. All crRNAs and HDR templates used in this study are listed in Table S1.

### Cell viability

TNBC cell lines including human MDA-MB-231 and mouse 4T1 cells were seeded into 96-well plate (M0812, Greiner CELLSTAR®, Germany) at the density of 5×10^3^ cells/ well. The cell viability was then evaluated 48 hours after IFN-exposure by using AlamarBlue™ (A50101, Thermo Scientific). The fluorescence was measured using plate reader (Spark 20M, TECAN, Switzerland) following the setting of 560 nm excitation and 590 nm emission wavelengths. The percentage of cell viability was calculated and normalised to control.

### Western blot analysis

At the end of experiment, cells were washed twice with cold PBS before extracting protein by using with RIPA lysis buffer containing 1X protease inhibitor cocktails (11836170001, Roche) and 1X phosphatase inhibitor (PhosSTOP; 4906845001, Roche). The protein concentration was measured by using Pierce™ BCA Protein Assay Kits (23227, Thermo Fisher)

Twenty-five micrograms of protein samples were resolved by sodium dodecyl sulphate polyacrylamide gel electrophoresis (SDS-PAGE) before being transferred to nitrocellulose blotting membrane (1620115, Bio-rad, CA, USA). The primary antibodies were incubated at 4 °C overnight. Primary antibodies used were STAT1 (14994S, Cell signaling), p-STAT1 (7649S, Cell signaling), STAT2 (MA5-42463, Invitrogen), p-STAT2 (88410S, Cell signaling), USP18 (4813T, Cell signaling), ISG15 (15981-1-AP, ProteinTech), MHC-I (MA5-35712, Invitrogen) and β-actin (A2066-100UL, Sigma Aldrich). After washing, the membranes were then incubated with the HRP-linked secondary antibody (7074S, Cell Signaling). Immunoblot membranes were probed with enhanced chemiluminescence (ECL) (Bio-rad) and visualized protein bands by using ChemiDoc MP (Bio-rad). Protein band intensity was quantified using ImageJ software (NIH). To relatively quantify the target protein expression, the intensity of target protein band was normalized to β-actin of particular lane.

### Immunofluorescence staining of cell surface marker

Cells were seeded onto 4-well culture slides (354114, Corning, USA) at a density of 2 × 10⁴ cells/well and allowed to adhere overnight. Cells were treated with 10 U/mL IFN-α2b for 48 hours. To preserve the native structure of cell surface proteins, cells were fixed in freshly prepared 4% paraformaldehyde (PFA) in PBS for 10 min at room temperature without permeabilization. Following fixation, cells were washed three times with PBS and incubated in blocking buffer (5% BSA in PBS) for 30 min. Primary antibodies (1:100) against MHC-I surface marker was applied to cells overnight at 4°C. After three washes in PBST, cells were incubated with Alexa Fluor™ 594-conjugated anti-rabbit IgG (H+L) secondary antibody (1:200) for 2 hours at room temperature in the dark. Slides were washed twice with PBST and mounted using antifade mounting medium containing DAPI (H-1200, VECTASHIELD®, USA). Protein expression was visualized using a Nikon AX confocal microscope (Nikon).

### Proteomic sample preparation

#### In vitro cell culture sample

MDA-MB-231 or 4T1 cells were seeded into 96-well plate at 5×10^3^ cells/well overnight before applying IFN-α2. After 48 hours of treatment, cells were wash twice with PBS followed by adding lysis buffer containing 1% (w/v) sodium deoxycholate (D6750, Sigma-Aldrich), 100 mM Tris pH 8.5 (T5941, Sigma-Aldrich), 1 mg/mL chloroacetamide (C0267, Sigma-Aldrich) and 1.5 mg/mL TCEP (C4706, Sigma-Aldrich). Samples were then heated at 95°C for 30 minutes before applying 5 *µ*g MS-grade trypsin (90058, Thermo Scientific) into each sample. All samples were followed by 37°C incubation overnight.

One volume of stage tip loading buffer containing 2% (v/v) TFA in isopropanol was added into each well at the end of incubation to acidify peptides. Then, peptides-containing solution were transferred to double disks styrene-divinylbenzene reverse phase sulfonate (SDB-RPS) stage tips for desalting process as described previously^38^. Briefly, peptides-containing stage tips were washed twice with stage-tip wash buffer A (1% (v/v) TFA in isopropanol) and stage-tip wash buffer B (0.2% (v/v) TFA in 5% (v/v) ACN), respectively. Peptides were then eluted by adding elution buffer, 1% (v/v) NH_4_OH in 60% (v/v) ACN, followed by drying using speed vacuum (Labconco, MO, USA). The tryptic peptides were reconstituted in 0.1% (v/v) TFA in MS-grade water.

#### In vivo tumour sample

4T1 tumour tissues were lysed in a buffer containing 5% (w/v) sodium dodecyl sulphate (SDS; L4509, Sigma-Aldrich), 100 mM Tris-HCl pH 8.5 (Sigma-Aldrich), 1 mg/mL chloroacetamide (Sigma-Aldrich), and 1.5 mg/mL TCEP (Sigma-Aldrich). The lysates were subjected to sonication followed by heat at 95°C for 20 minutes to denature proteins and reduce disulphide bonds. Protein aggregation capture (PAC) was then performed using MagReSyn® HILIC beads (MR-HLC002, ReSyn Biosciences) on a KingFisher™ Flex purification system, as described previously^39^. Tryptic digestion was carried out, and resulting peptides were acidified with 2% (v/v) formic acid. Peptides were desalted using C18 stage tips. Briefly, stage tips were first equilibrated with absolute methanol followed by C18 stage tip wash buffer consisting of 0.1% (v/v) trifluoroacetic acid (TFA) in MS-grade water. Peptides were then eluted using 0.1% (v/v) TFA in 50% (v/v) ACN followed by drying in a speed vacuum concentrator (Labconco). The peptides were reconstituted in 0.1% (v/v) TFA in MS-grade water for further LC-MS/MS analysis.

To study proteomic profile, all samples were injected into a Orbitrap Fusion Lumos mass spectrometer (Thermo Scientific) which was operated in data independent acquisition (DIA) mode.

### LC-MS setting

Desalted peptides were then loaded onto 25cm Aurora Columns (IonOptiks, Australia) using a RSLC nano µHPLC system connected to a Fusion Lumos mass spectrometer. Peptides were separated by a 70 min linear gradient from 5% to 30% acetonitrile, 0.5% acetic acid. The mass spectrometer was operated in DIA mode, acquiring a MS 350-1650 Da at 120k resolution, followed by MS/MS on 45 windows with 0.5 Da overlap (200-2000 Da) at 30 k with a NCE setting of 28.

For proteomic analysis, raw files were analysed by using DIA-NN 2.0 (https://github.com/vdemichev/DiaNN) for searching against the UniProt *Homo sapiens* database for MDA-MB-231 samples and *Mus musculus* for 4T1 cells. The deep-learning option was used for obtaining the precursor ion spectra from the FASTA files with the default setting for exception of “Robust LC (high precision)”.

The proteomics data including raw files and search results have been deposited to the ProteomeXchange Consortium via the PRIDE partner repository with the dataset identifier PXD064305, PXD064246, PXD064445, PDX064479 and PDX064517.

### USP18 structural analysis

To investigate the impact of arginine substitutions, Alpha Fold was used to analyse to potential effect ^40^. The predicted structure between USP18^WT^ and USP18^C64R/C65R^ was then superposed to compare structural differences using Chimera X1.9 ^41^.

### USP18-STAT2 mediated type I IFN suppression study

To further investigate the impact of USP18 inhibition and its disruption on STAT2-dependent IFN-response inhibition, MDA-MB-231 cells were primed with 250 U/mL IFN-α2b for 6 hours. Thereafter, treatment was removed followed by PBS washes. Cells were rested for 18 hours. Then, 250 U/mL IFN-α2b was applied to pulse the IFN-response in MDA-MB-231 cells for 1 hour. Proteins were collected accordingly by using phosphatase-/ protease-inhibitor containing RIPA buffer extraction. Twenty-five micrograms of protein were loaded and the IFN-related signature was analysed by western blot analysis.

### Cytotoxic assay of NK-92 cells on MDA-MB-231 in co-culture system

To investigate the cytotoxic effect mediating apoptosis of MDA-MB-231 cells in NK-92 coculture system, MDA-MB-231 cells were pre-labelled with 1 *µ*M CellTracker™ Orange CMRA Dye (C34565, Invitrogen, MA, USA) for 30 minutes at 37°C. Pre-labelled MDA-MB-231 cells were then seeded into 96-well plate (Greiner CELLSTAR®). Twenty-four hours after seeding, cells were treated with 10 U/mL IFN-α2b (pbl assay science) for 24 hours. Thereafter, culture medium was replaced with cytotoxic detection medium, which contains NK-92 cells and 1 *µ*g/mL CellEvent™ Caspase-3/7 (Invitrogen). The cytotoxic effect NK-92 on MDA-MB-231 cells was monitored using Incucyte S3 instrument (Sartorius) and measured fluorescent puncta by using ImageJ software.

### Mice

All animal research was approved by the University of Edinburgh Animal and Ethical Review Board (PL05-21) and carried out under UK Home Office Project licence PP7510272. Female BALB/c mice were 6-12 weeks old at the start of the experiment. Mouse TNBC cells (4T1) were subcutaneously injected into both flanks at a concentration of 5×10^4^ cells in 100 *µ*L serum-free RPMI-media. Tumour growth was observed and measured biweekly using callipers. For survival studies mice were sacrificed when tumours (unilateral) reached 15 mm in diameter, as defined in PP7510272. Tumours were also collected at 20 days post implantation for immune profiling, and snap-frozen for proteomic analysis.

For ICIs treatment study, 6-12 weeks old mice were subcutaneously implanted 4T1 cells. *InVivoMAb* anti-mouse PD-1 (clone RMP1-14; BE0146, BioXcell) was intraperitoneally injected at 250 *µ*g in PBS on day 3, 7, 10, 14, and 17 after implantation. Tumour size was monitored by using callipers biweekly and tumours collected on Day 20.

### Immune cell profiling analysis

Tumour samples were digested with Liberase TL (5401020001, Roche) and DNase I (69182-3, Sigma-Aldrich) for 30 minutes at 37°C in a shaking incubator. The digested tissues were then passed through a 70 *µ*m cell strainer (10788201, Falcon). Red blood cells were subsequently removed from all sample by using RBC Lysis Buffer (420301, BioLegend).

Following tissue dissociations, cells were stained with LIVE/DEAD™ Fixable Blue (L23105, Invitrogen) and resuspended in PBS + 1% BSA. Approximately 1 × 10⁶ cells were aliquoted into 5 mL tubes and incubated with Fc Blocking antibody (101319, BioLegend). Antibodies used for immune profiling are listed in Table S2, and master mixes were prepared prior to incubation with cells. After washing, cells were fixed and permeabilized overnight with eBioscience™ FOXP3/transcription factor staining buffer set (00-5523-00, Invitrogen) following the manufacturer’s instructions. Cells were then stained with intracellular antibody master mixes, washed, and acquired on a BD LSR Fortessa (BD Biosciences). Flow cytometry data analysis and gating were performed using FlowJo v10.10 (Tree Star).

### Statistical analysis

All results were statistically analysed and visualised using GraphPad Prism 10 (GraphPad Software Inc., USA). One-way or two-way analysis of variance (ANOVA) and multiple comparisons Tukey post-test were used to examine statistical different between groups. While student T-test was used to compare the difference of mean between 2 groups. P-values less than 0.05 were regarded as statistically significant for all statistical analyses.

## Results

### USP18 is the essential regulator of type I IFN-response in TNBC

To investigate the expression of USP18 in TNBC, we conducted a meta-analysis using data from TCGA and GTEx. We found that USP18 expression was significantly upregulated in breast cancer compared to normal tissues. (fig. S1A-B). Furthermore, the USP18 expression was commonly upregulated across breast cancer subtypes, including TNBC (fig. S1C). As TNBC is one of the most aggressive subtypes and exhibits a cold tumour immune response, we aimed to investigate how USP18 loss affected the IFN response in MDA-MB-231, a human TNBC cell line. We assayed cells after 48 h and found that loss of USP18 significantly decreased viability upon IFN-α2b treatment (fig. 1A). A similar effect was also seen in mouse TNBC (4T1) cells, as shown in fig. S2A. The reduction of cell viability demonstrated that USP18 loss increased IFN sensitivity in both cell lines. To elucidate the molecular changes, proteomic analysis was carried out. MDA-MB-231 cells were treated with 10 U/mL IFN-α2b for 48 h, followed by proteomic analysis, which revealed significant alterations in the proteome as illustrated in a volcano plot (fig. 1B). The significantly changed proteins were analysed by gene ontology enrichment for biological process (GOBP). The type I IFN-response, innate immune response and cytokine production were all upregulated (fig. 1C) as well as the ISG-protein signature (fig. S2B). We further identified the downregulation of cell proliferation-related processes such as the cell cycle, cell division, and microtubule cytoskeleton organisation (fig. 1D). Our findings confirmed that, in our cell models, USP18 is an important mediator and negative regulator of the cellular IFN response.

**Figure 1.**
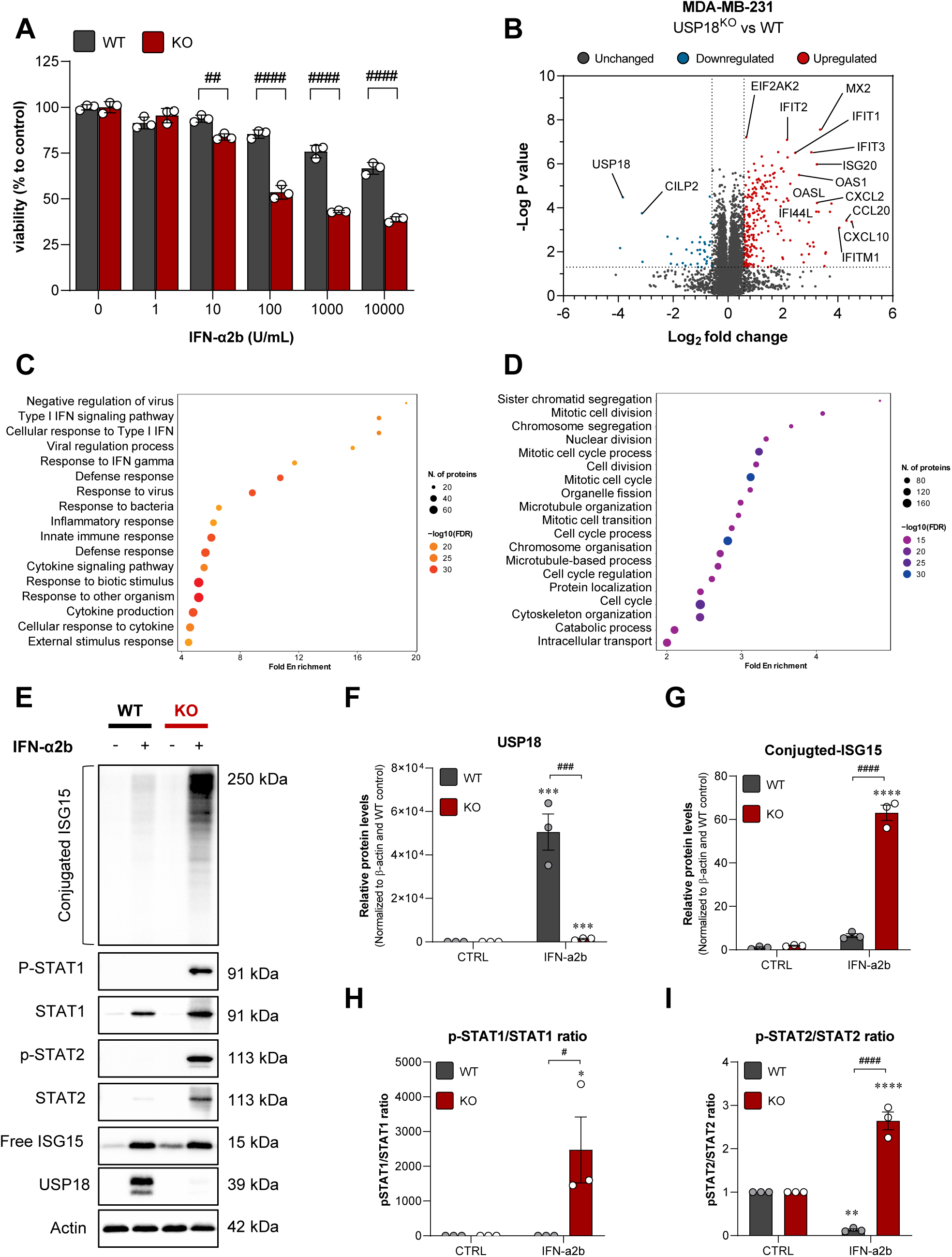
USP18 plays an important role in IFN susceptibility in TNBC. **(A)** Cell viability of IFN-α2b on USP18^WT^ and USP18^KO^. **(B)** A volcano plot of proteome alteration in USP18^KO^ vs USP18^WT^ after 48 h of 10 U/mL IFN-α2b treatment. Enrichment analysis of **(C)** upregulated proteins and **(D)** downregulated proteins. **(E)** Immunoblot of MDA-MB-231. The quantification of immunoblots were shown as **(F)** USP18, (**G)** conjugate-ISG15, **(H)** p-STAT1/STAT1 and **(I)** p-STAT2/STAT2 ratio. Graphs showed mean of 3 independent biological replicates ± SEM. *p<0.05, ***p<0.001, and ****p<0.0001 versus control, while #p<0.05, ###p<0.001, and ####p<0.0001.

To further validate the type I IFN-sensitive phenotype of MDA-MB-231 cell lysates were collected for immunoblot analysis. We observed that USP18 expression was significantly upregulated in USP18^WT^ cells following IFN-α2b stimulation, confirming that USP18 protein expression is induced by the type I IFN signalling pathway (fig. 1E,F). In addition, both conjugated ISG15 and free ISG15 levels were found to be significantly elevated following IFN-α2b stimulation, with a markedly stronger induction observed in USP18^KO^ cells (fig. 1E,G and S2G). The results also showed a significant increase in p-STAT1, p-STAT2, STAT1, and STAT2 levels in USP18^KO^ MDA-MB-231 cells following IFN-α2b exposure (fig. 1E, S2C-F). Furthermore, p-STAT1/STAT1 and p-STAT2/STAT2 ratio were significantly increased in USP18^KO^ MDA-MB-231 cells (figs. 1E, H, I). This increase indicates an upregulation of IFN response signalling and ISG expression in USP18^KO^ MDA-MB-231 cells, whereas the response was dampened in USP18^WT^ MDA-MB-231 cells, consistent with USP18’s role as a negative regulator of IFN signalling.

### Loss of USP18 induces immune-response related proteins in TNBC

Given that type I IFN signalling plays a role in mediating immunity during infection, we further examined the impact of USP18 loss on the immune response. Gene set enrichment analysis (GSEA) was performed on the proteomic data, revealing a positive correlation of innate immune response and adaptive immune response in USP18^KO^ MDA-MB-231 cells (fig. S3A-B, D-E). Moreover, we observed significant GO enrichment in selected cellular components (GOCC) in USP18^KO^ cells following IFN exposure (fig. 2A). The GOCC human leukocyte antigen (HLA) family was most significantly upregulated, suggesting a broad induction of the major histocompatibility complex (MHC) proteins in the absence of USP18 (fig. 2B and S3C,F), indicating enhanced antigen presentation and immunogenicity. To further investigate, immunoblot and immunofluorescence analyses were performed to assess MHC-I expression in MDA-MB-231 cells. The results showed a significant increase in MHC-I levels in USP18^KO^ cells following IFN-α2b exposure (fig. 2C-D). Furthermore, Fas cell surface death receptor (Fas), which plays an important role in activating apoptosis cell death through FAS/ FASL interactions, was also significantly increased in USP18^KO^ cells (fig. 2E). The upregulation of MHC-I and Fas by IFN-α2b exposure in cancer cells may improve immune recognition by immune cells such as NK cells and T lymphocytes highlighting a potentially stronger anti-tumour immune response in USP18^KO^ cells. As a result, these cells may become more vulnerable to NK/ T cell-mediated killing, p. To investigate the cytotoxic effect of immune cells mediating cancer cell death, a co-culture system between MDA-MB-231 and the NK cell line NK-92 was conducted. As shown above, USP18^KO^ MDA-MB-231 cell viability was significantly decreased under IFN-α2b treatment. Moreover, cell viability was further reduced by NK-92 co-culture in both cell lines, but more pronounced in the absence of USP18 (fig. 2F). Taken together, these data suggest that USP18 loss increases MHC, Fas expression and sensitises cells to NK-mediated cell death.

**Figure 2.**
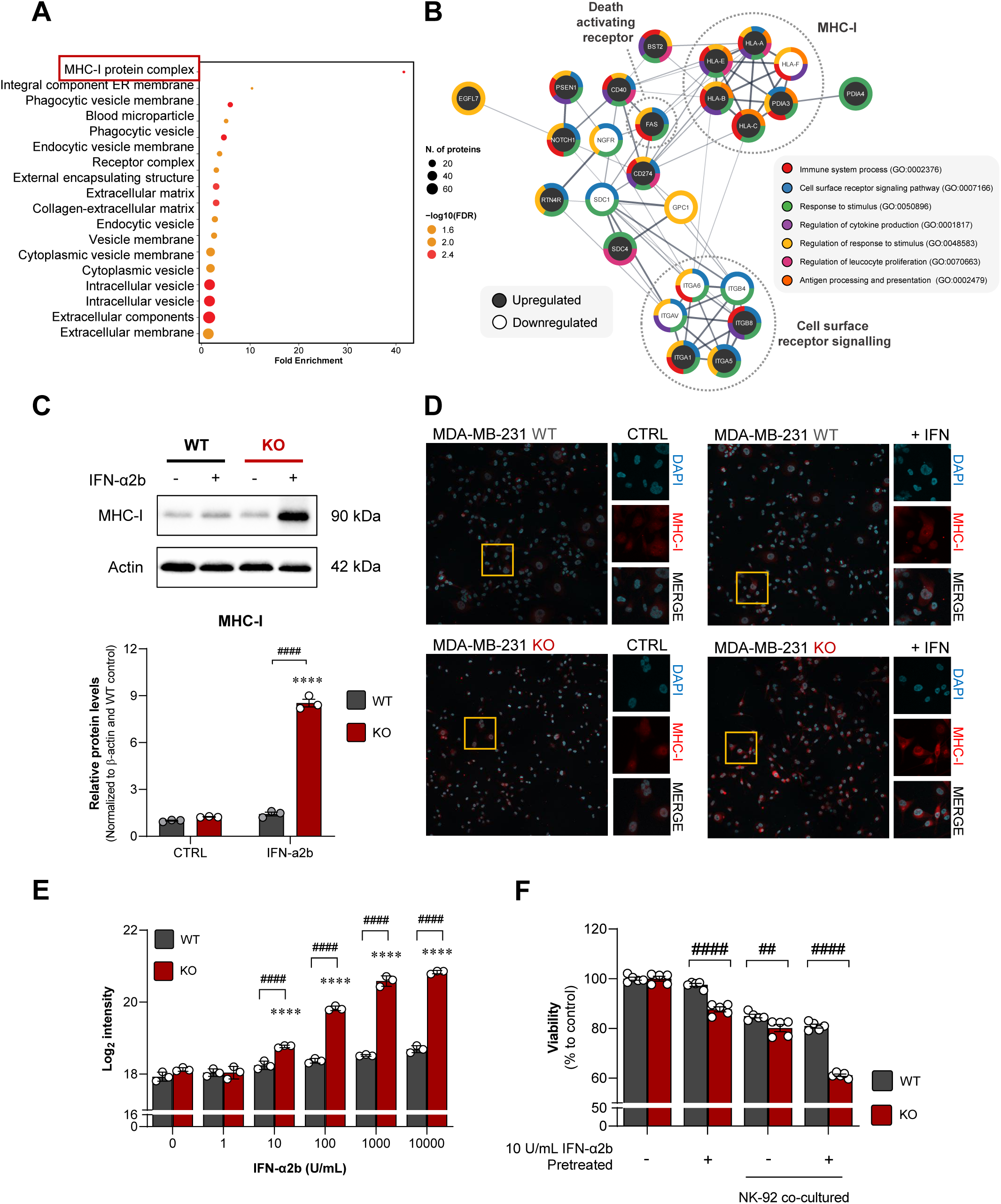
USP18 loss enhanced IFN-inducing immune response mediating immunogenic modulation in TNBC. **(A)** Gene ontology enrichment of cellular component proteomic alteration in USP18^KO^. **(B)** Protein-protein interaction diagram of cellular component related immune response. The expression analysis of MHC-I in MDA-MB-231 by **(C)** immunoblot and **(D)** immunofluorescent. **(E)** Fas expression. **(F)** Cytotoxic effect of NK-92 on MDA-MB-231 in co-culture system. Graphs showed mean of at least 3 independent biological replicates ± SEM. ****p<0.0001 versus WT control; while ##p<0.01 and ####p<0.0001.

### USP18 loss impacted tumour growth and the immune cell landscape in the TNBC TME

To determine whether loss of USP18 regulated tumorigenesis and the TME in an immune-competent murine model, USP18^KO^ and USP18^WT^ 4T1 cells were subcutaneously injected into immune competent BALB/c mice. Overall survival was only modestly increased by USP18 loss (fig. 3A). Nevertheless, tumour growth was significantly reduced, resulting in significantly smaller tumours at day 20 post-implantation (fig. 3B-C). To determine protein expression changes, tumours were collected at Day 20 and analysed by proteomics. As expected, USP18 levels were significantly decreased (fig. 3D) and immune activation-related pathways were significantly enriched in USP18^KO^ tumours, indicating an increased immune response (fig. 3E-F). Interestingly, we were unable to identify a proliferative signature, suggesting that the reduction in tumour mass may be driven by immune activation in USP18-deficient tumours.

**Figure 3.**
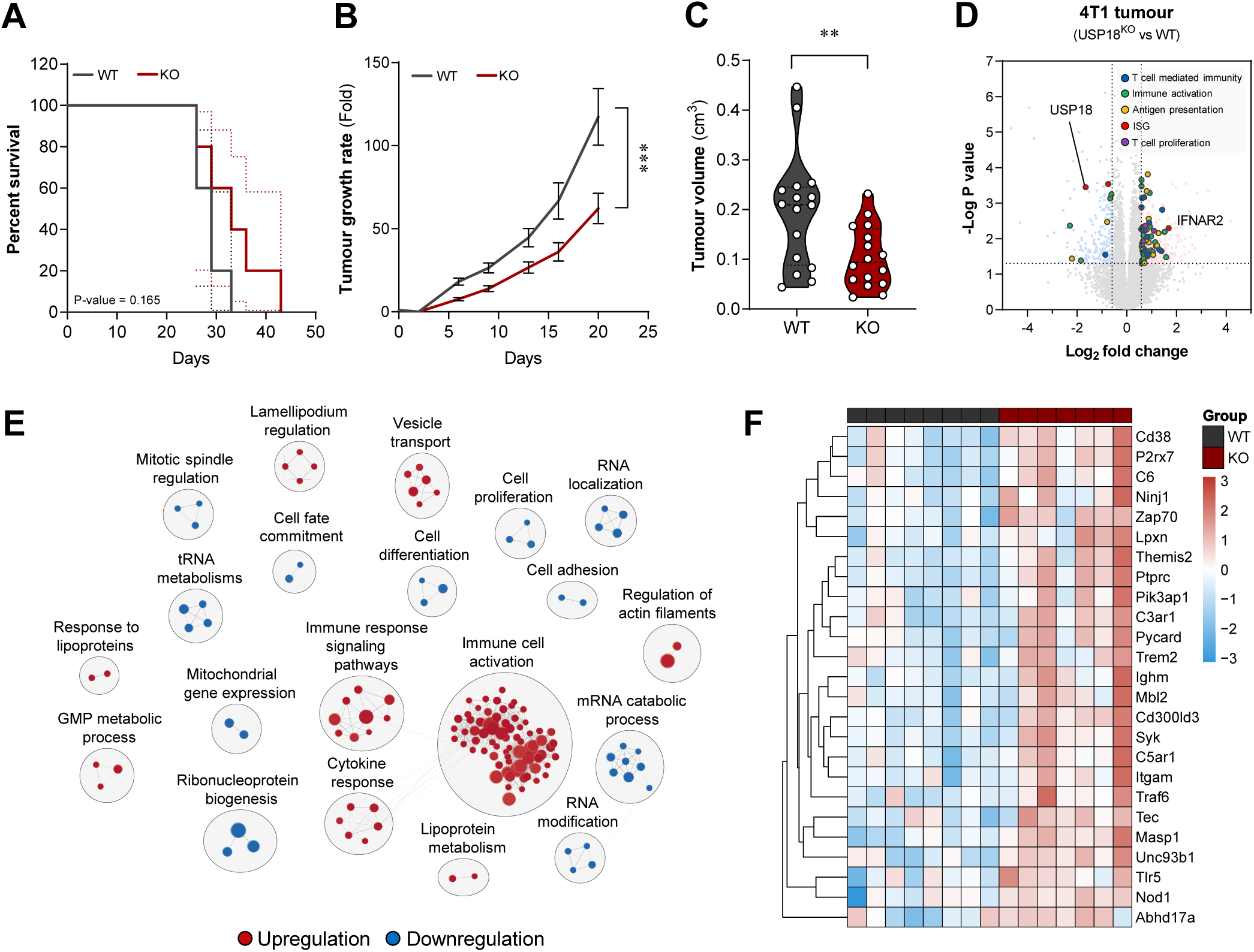
Loss USP18 reduced TNBC tumour growth and activated immune response *in vivo*. **(A)** Overall survival. **(B)** Tumour growth rate. **(C)** Tumour volume at day 20 after implantation. **(D)** Volcano plot of tumour proteomics. **(E)** Gene ontology enrichment of tumour proteomics. **(F)** Top 25 immune activation related protein signatures. Graphs showed mean size of 16 tumours ± SEM collected from 8 mice per group and 8 tumours per group for proteomic analysis. ***p<0.0001 versus WT tumours.

To further investigate how USP18 depletion shapes the breast cancer TME, we profiled tumour infiltrating immune cells using flow cytometry. As outlined in fig. S4–S5, we segmented the major immune cell lineages and observed a notable expansion in the overall immune compartment. Specifically, the number of CD45⁺ immune cells was significantly elevated in USP18^KO^ tumours compared to USP18^WT^ tumours (fig. 4A), pointing toward a more immune-enriched TME upon USP18 loss. Delving further, we uncovered differences in myeloid subtypes in USP18^KO^ tumours, including a significant reduction in CD11b⁺ myeloid cells (monocytes, macrophages, granulocytes, and dendritic cells) and a substantial decrease in neutrophil infiltration (fig. 4B). This was accompanied by an elevated presence of conventional dendritic cells type 1 (cDC1), whereas the cDC2 population remained largely unchanged (fig. 4C). Interestingly, the number of monocyte abundance in TME remain unchanged, although we detected a significant rise in monocyte-derived dendritic cells (moDCs) within the USP18^KO^ TME (fig. 4B–C), suggesting a shift in myeloid cell differentiation or recruitment. Furthermore, the F4/80^+^ macrophage population was significantly increased (fig. 4D). Notably, macrophage polarization appeared altered as well: the pro-inflammatory macrophage subset rose by 22.17%, while anti-inflammatory macrophages showed a modest 3.30% increase (fig. 4D), reflecting broader reprogramming of myeloid phenotypes in the absence of USP18. To gain further insight into the functional status of these myeloid cells, we assessed PD-L1 expression across the infiltrating myeloid subsets. The results revealed a reorganization of immune signalling. PD-L1 levels were modestly reduced in neutrophils and were significantly reduced in anti-inflammatory macrophages (fig. S6A–B), consistent with a less immunosuppressive microenvironment. In contrast, PD-L1 expression was elevated in pro-inflammatory macrophages, moDCs, and cDC2s (fig. S6C–E), suggesting an IFN-driven inflammatory program.

**Figure 4.**
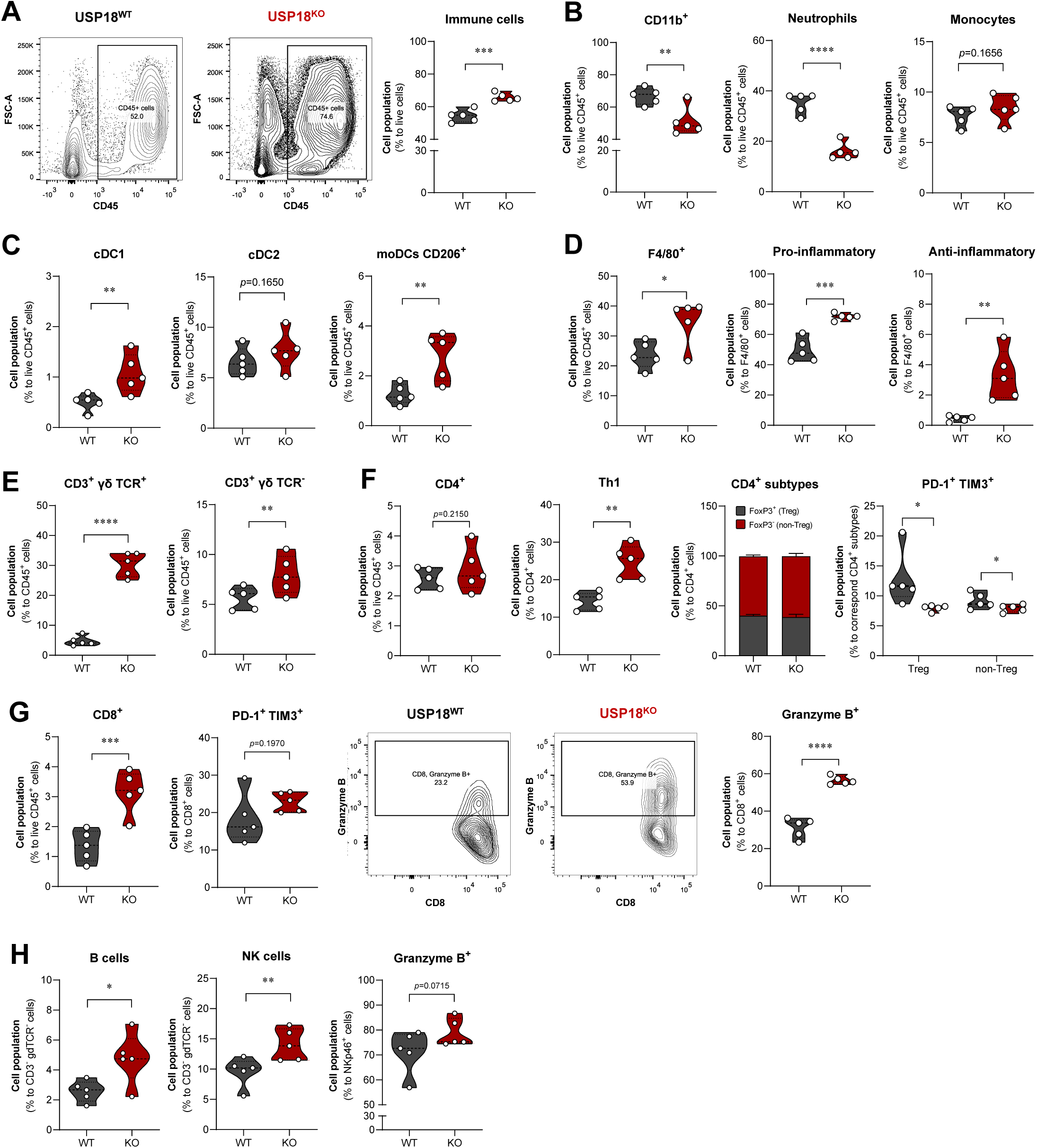
Loss of USP18 modulates immune cell population in the TME. Flow cytometry analysis showed **(A)** Proportion of immune cells in TME. **(B-D)** Myeloid cell subtypes in in tumours. **(E-G)** CD3^+^ T cell subtypes in tumours. **(H)** CD3^-^ immune subtypes. Graphs showed percentage of cell population from 5 mice ± SEM (5 tumours per group). *p<0.05, **p<0.01, ***p<0.001, and ****p<0.0001 versus WT tumours.

Given the central role of myeloid cells in orchestrating T cell responses, we then investigated T cell dynamics in the TME. USP18^KO^ tumours exhibited a pronounced increase in both CD3⁺ γδ TCR⁺ and CD3⁺ γδ TCR⁻ populations (fig. 4E), indicating an overall expansion of T cells within the TME. Among the T cell subsets, CD4⁺ T cells showed a modest increase in USP18^KO^ tumours (fig. 4F), while there was a significant increase in T helper 1 (Th1) cells (fig. 4F). Although the loss of USP18 did not alter the overall number of Treg cells, it was associated with an increased frequency of exhausted CD4⁺ T cells co-expressing PD-1 and TIM-3, suggesting that USP18 deficiency exacerbates T cell dysfunction (fig. 4F).

The observed increase in cDC1s—key activators of cytotoxic CD8⁺ T cells—led us to examine the CD8⁺ T cell compartment. Consistent with this, USP18^KO^ tumours displayed a marked enrichment of CD8⁺ T cells, with a 15.05% increase compared to USP18^WT^ tumours (fig. 4G). Despite this expansion, we did not observe any significant changes in the frequency of exhausted CD8⁺ T cells, as defined by co-expression of PD-1 and TIM3. However, the proportion of granzyme B⁺ CD8⁺ T cells was elevated by 25.15%, highlighting a shift toward a more activated and cytotoxic T cell phenotype (fig. 4G).

Beyond T cells, the landscape of lymphocytes also shifted. We found a notable increase in B220⁺ B cells, NKp46⁺ natural killer (NK) cells, and granzyme B⁺ NK cells in the USP18^KO^ TME (fig. 4H). These changes collectively point to a more inflamed, or “hot,” immune contexture in the 4T1 tumours.

Lastly, we evaluated the therapeutic implications of this immunological reprogramming. In line with a more immunostimulatory TME, USP18 deficiency markedly enhanced the efficacy of anti–PD-1 (αPD-1) therapy. Treated USP18^KO^ tumours displayed reduced growth and a heightened immune response (Fig. S7), reinforcing the role of USP18 as a key immunoregulatory node in breast cancer. Altogether, these findings reveal that USP18 loss profoundly reshapes the immune landscape of the TME—enhancing both innate and adaptive immune components—and sensitises TNBC tumours to immune checkpoint blockade.

### USP18 inhibition mimicry induces IFN-sensitive phenotype in TNBC

Given our observation that USP18 deletion enhances IFN signalling and increases immunogenicity in TNBC cells and tumours, we sought to determine whether mimicking USP18 inhibition by an inhibitor would yield a similar effect. To simulate the impact of a small-molecule inhibitor, we used CRISPR-Cas9 to introduce arginine (R) substitutions at the catalytic cysteine residues (C64 and C65) of USP18 in MDA-MB-231 cells (fig. 5A). Consistent with previous experiments, MDA-MB-231 cell viability was significantly reduced upon IFN-α2b treatments in a concentration-dependent manner, with the most pronounced effect observed in USP18^KO^ and USP18^C64R/C65R^ clones (fig. 5B). This indicated that mutating USP18 effectively increase IFN sensitivity by enhancing the IFN signalling pathway.

**Figure 5.**
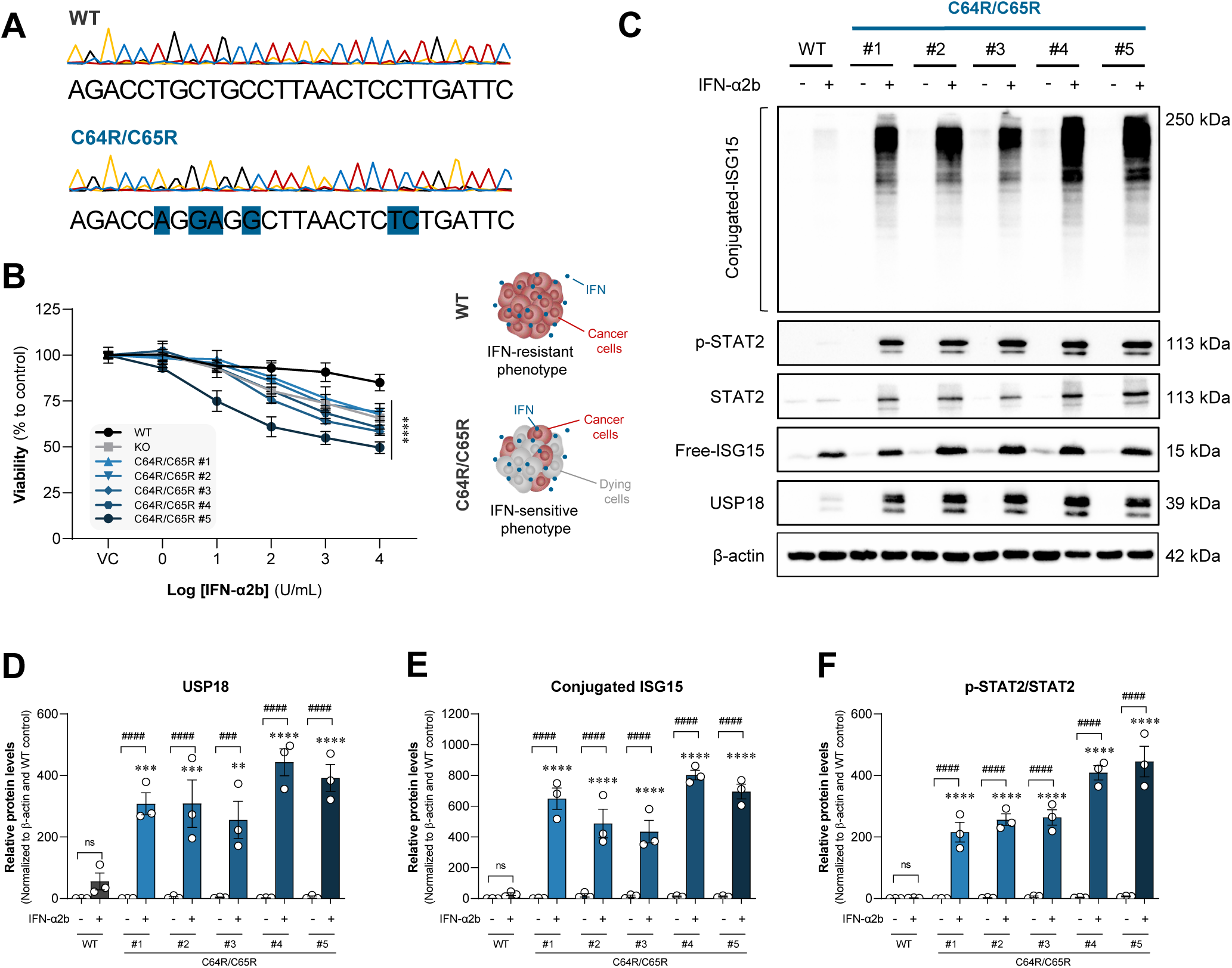
Inhibition of USP18 functions enhanced IFN response in TNBC. **(A)** DNA sequencing result representing C64R/C65R substitution of USP18. **(B)** Cell viability of USP18^C64R/C65R^ MDA-MB-231 clones after 48h of IFN-α2b treatment (n=6). **(C)** Immunoblot represented level of downstream signalling of type I IFN response after 48h of IFN-α2b treatment (n=3). The quantification of **(D)** USP18; **(E)** conjugated-ISG15; and **(F)** p-STAT2/STAT2 ratio. Graphs showed mean of at least 3 independent biological replicates ± SEM. **p<0.01, ***p<0.001, and ****p<0.0001 versus WT control. While ###p<0.001, and ####p<0.0001.

To functionally characterise the effects and molecular consequences of C64R/C65R mutation in TNBC, an IFN stimulation experiment was conducted by treating MDA-MB-231 cells with 10 U/mL IFN-α2b for 48 hours. USP18 expression was significantly elevated in USP18^C64R/C65R^ cells, indicating an enhanced interferon (IFN) response and the cells’ attempt to counteract IFN response (Fig. 5C-D). In addition, ISG15 levels were notably increased in USP18^C64R/C65R^ cells, consistent with an enhanced IFN-response (fig. 5C, E and S8A). We also observed significantly increased levels of p-STAT2, STAT2 and p-STAT2/STAT2 in USP18^C64R/C65R^ cells (fig. 5C,F and S8C-D), indicating that mutant USP18 phenocopied what we observed in the KO lines.

### Modification of USP18’s active site induced subtle structural alterations

Previous studies have indicated that the amino acid region 51–112 of USP18 plays a critical role in STAT2-dependent binding, which mediates IFN response suppression ^29,42^. To further identify the effect of C64R/C65R substitution, we predicted the structure of the WT and mutant USP18 using Alphafold V. 3. The introduction of the larger and charged side chains of R into the active site of USP18 induced changes of the catalytic cleft loop structure (fig. 6A). This structural alteration may disrupt STAT2 binding, potentially explaining the increased IFN sensitivity typically linked to USP18’s non-enzymatic function. By interfering with the STAT2–USP18 interaction, the mutation could impair the negative feedback loop that relies on USP18’s scaffolding role.

**Figure 6.**
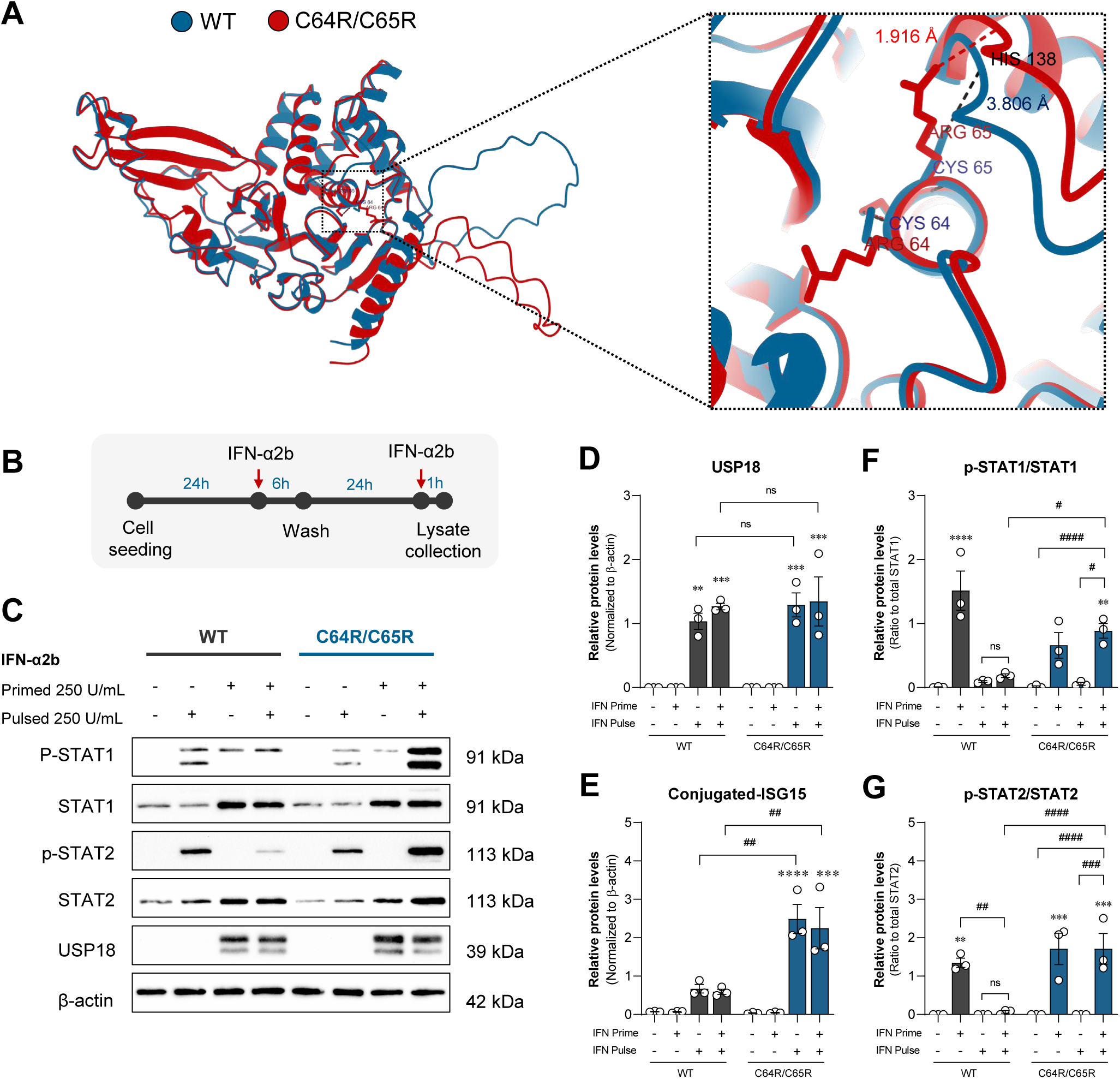
Catalytically mutated USP18 prolonged IFN response in MDA-MB-231 cell by STAT2-binding disruption. **(A)** Alpha fold analysis of hUSP18^C64R/C65R^ versus hUSP18^WT^. **(B)** Timeline of time course experiment investigating the suppression of negative regulation role of USP18. **(C)** Immunoblot represented level of downstream signalling of type I IFN response from time course experiment. The quantification of **(D)** USP18; **(E)** conjugated-ISG15; **(F)** p-STAT1/STAT1 ratio and **(G)** p-STAT2/STAT2 ratio. Bar graphs showed mean of 3 independent biological replicates ± SEM. **p<0.01 and ***p<0.001 versus WT control. While #p<0.05, ##p<0.01, ###p<0.001, and ####p<0.0001.

To determine whether the active site mutation disrupted the non-enzymatic feedback inhibitory loop, we performed a prime and stimulate experiment (fig. 6B). Cells were first primed with 250 U/mL IFN-α2b for 6 hours, followed by a 24-hour resting phase. Subsequently, cells were exposed to a 1-hour pulse of 250 U/mL IFN-α2b. We observed induction of the IFN response following the initial priming treatment in both USP18^WT^ and USP18^C64R/C65R^. While no significant differences were detected in the expression of interferon-stimulated genes (ISGs) such as USP18 and ISG15 between USP18^WT^ and USP18^C64R/C65R^ cells after the prime + pulse treatment (fig. 6V and S9), phosphorylated STAT1 and STAT2 (p-STAT1 and p-STAT2) levels were markedly elevated in USP18^C64R/C65R^ cells but not in USP18^WT^ cells. This led to increased p-STAT1/total STAT1 and p-STAT2/total STAT2 ratios in the mutant cells (fig. 6C, F–G, and S9).

These findings suggest that the C64R/C65R mutation prevents effective negative feedback regulation of type I IFN signalling by USP18. Consequently, targeting USP18’s active site with an inhibitor could be a strategy to inhibit both the enzymatic and scaffolding functions of USP18 and sensitise cell to IFN.

### Catalytically mutated USP18 enhanced cancer immunogenicity

To determine the molecular phenotype of the IFN-sensitivity of cells harbouring the C64R/C65R mutation, we analysed the proteome of the cell line in the presence and absence of IFN. The results indicated that C64R/C65R mutation enhanced both the type I IFN response and the innate immune response in USP18^C64R/C65R^ MDA-MB-231 cells and USP18^C61R/C62R^ 4T1 cells (fig. 7A-B and S10A-C). We also observed a pronounced upregulation of MHC-I in USP18^C64R/C65R^ cells following IFN-α2 b treatment (Fig. S 10 D), which we confirmed by immunoblotting and immunofluorescence (fig. 7C-D). In addition, we observed an upregulation of the FAS protein (fig. S10E). To determine whether mutated USP18 increased FAS-driven cell death, we performed a co-culture experiment. We found that IFN-α2b treatment together with NK-92 co-culture significantly increased apoptosis in MDA-MB-231 harbouring USP18^C61R/C62R^ (fig. 7 E-F). Overall, these data suggest that USP18^C61R/C62R^ phenocopies the knockout and may increase cellular immunogenicity driven by IFN sensitivity.

**Figure 7.**
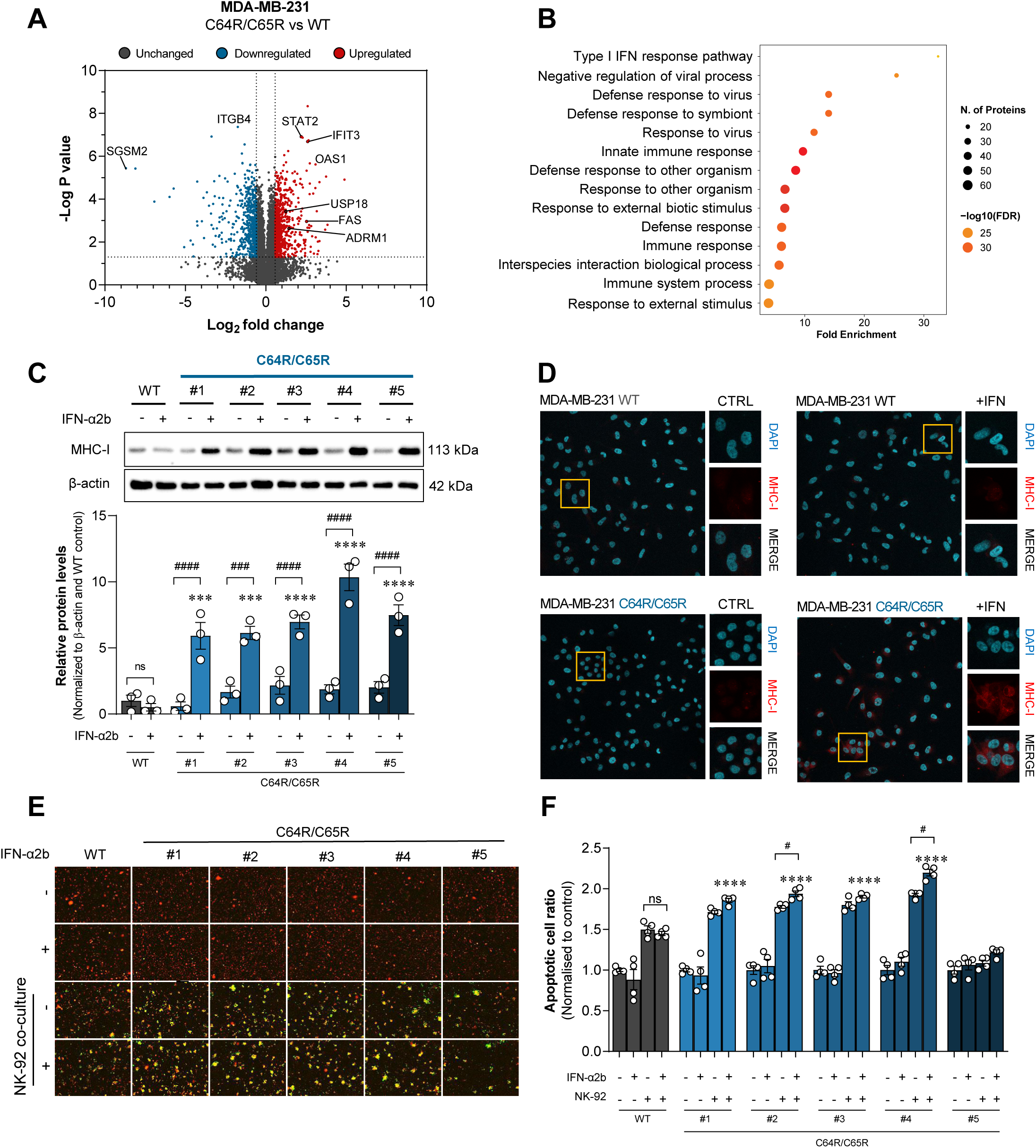
Catalytically mutated USP18 promoted IFN response inducing breast cancer immunogenicity. **(A)** Volcano plot of proteome alteration in USP18^C64R/C65R^ vs USP18^WT^ after 48 h of 10 U/mL IFN-α2b treatment. **(B)** Gene enrichment analysis. **(C)** Immunofluorescent MHC-I cell surface expression. **(D)** Immunofluorescent MHC-I cell surface expression. **(E-F)** Cytotoxic effect mediating apoptosis of MDA-MB-231 in NK-92 co-culture system. Bar graphs showed mean of at least 3 independent biological replicates ± SEM. ***p<0.001, and ****p<0.0001 versus control. While #p<0.05, ### p<0.001, and #### p<0.000.

### USP18^C61R/C62R^ reduced tumour growth *in vivo* by promoting immune-responsive TME

To further investigate the effects of USP18 and its disruption, 4T1 cells were subcutaneously implanted into BALB/c mice. We observed a significantly slower growth rate of tumour in USP18^KO^ and USP18^C61R/C62R^ compared to USP18^WT^ tumours (fig. 8A-B). To determine the mechanism underpinning the phenotype, we analysed the proteome of the tumours (fig. 8C). The activation of the immune response, B cell differentiation, myeloid activation, cytokine production and leukocyte intravasation were significantly upregulated in USP18^C61R/C62R^ tumours (fig. 8D-E). In addition, immune activation-related pathways were shown to be positively correlated with USP18^C61R/C62R^, including the innate immune response, adaptive immune response, and antigen presentation (fig. S11). These data suggested that mutation of USP18 augments the IFN-response, induces immune activation, and likely limits tumour growth.

**Figure 8.**
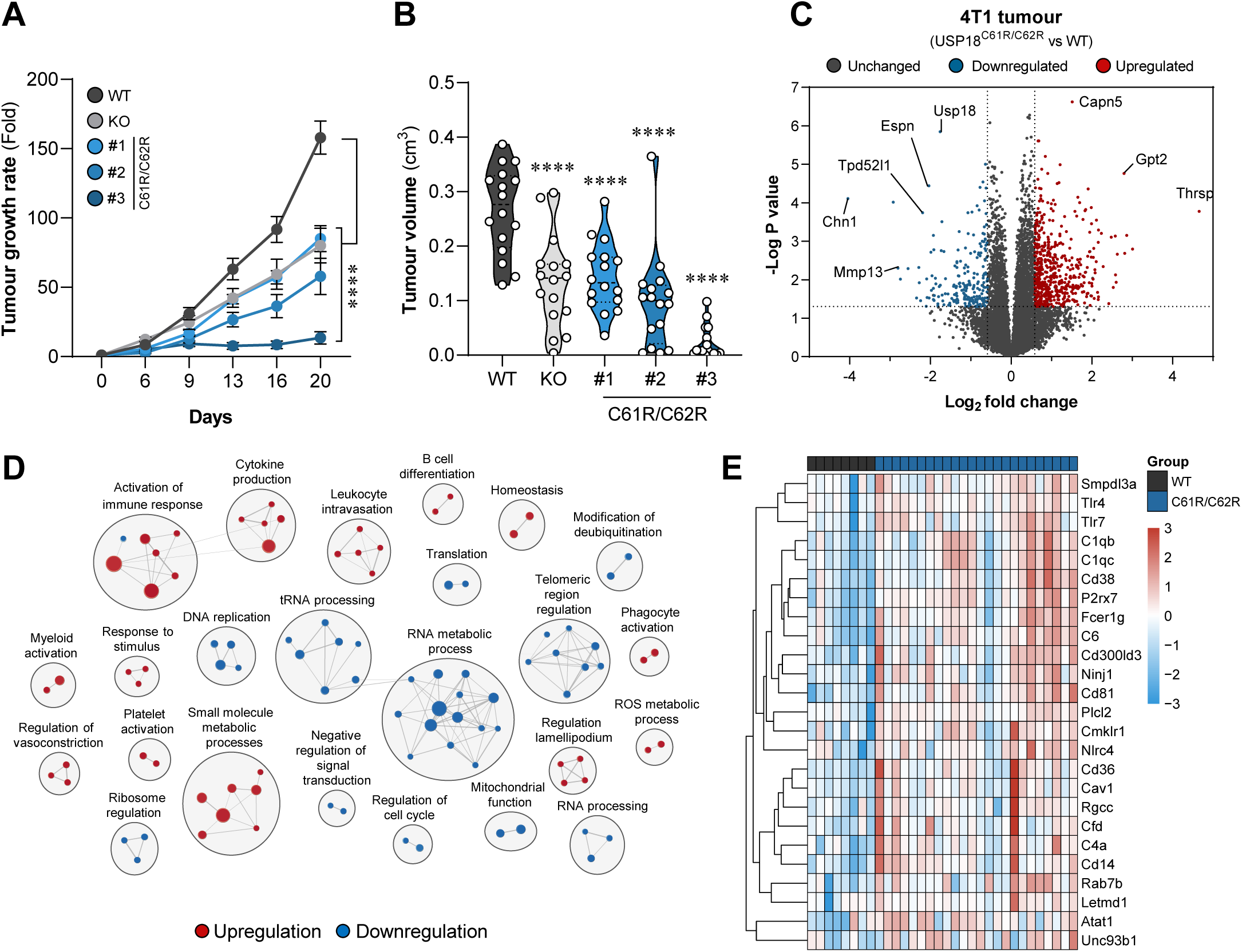
USP18 inhibition reduced tumour growth and activated immune response in the TME. **(A)** Tumour growth rate. **(B)** Tumour volume at day 20 after implantation. Tumour proteomic study was utilised showing; **(C)** Volcano plot, **(D)** Gene set enrichment, **(E)** Top 25 immune activation related protein signature. Graphs showed mean size of 16 tumours ± SEM collected from 8 mice per group. ****p<0.0001 versus WT tumours.

Our findings revealed the significant role of USP18 in cancer immunogenicity and immune response, and suggest that inhibition of USP18 with a small-molecule active-site binder could be a viable strategy to promote tumour immunogenicity in TNBC.

## Discussion

Type I interferons (IFNs) play a crucial role in anti-tumour immunity by activating cytokine signalling, stimulating immune responses, and promoting cell death ^20,21^ with USP18 as one of the key negative regulators of the pathway. Previous studies have shown that USP18 disrupts the formation of the IFNAR receptor complex, thereby dampening IFN responses in cancer cells ^26,29,33^. As a result, elevated USP18 expression can impair immune activation within the TME ^43^.

In this study, we demonstrated that loss of USP18 results in an IFN-hypersensitive phenotype, which enhanced cancer immunogenicity. Proteomic profiling revealed upregulation of surface proteins, such as MHC-I and Fas. Since MHC-I molecules are essential for antigen presentation and immune activation ^44–46^, their increased expression suggests improved immunogenic potential of TNBC cells. Prior studies have also reported a positive correlation between MHC-I expression and immunogenic cancer phenotypes ^47,48^. Additionally, Fas-a death receptor involved in apoptotic signalling ^49–51^-was significantly upregulated in USP18-depleted cells. These findings suggest that USP18 loss may improve tumour immunogenicity and sensitise cancer cells to immune-mediated killing processes. Crucially, we observed an increase in tumour immunogenicity *in vivo*. We observed a significant increase in overall immune cell infiltration in USP18-deficient tumours, indicating improved immune engagement within the TME. Further analysis revealed that USP18 loss was associated with changes in the composition of the immune cell landscape. Given that distinct immune cell subtypes exert divergent effects on tumour progression—such as pro-tumorigenic populations (e.g., anti-inflammatory macrophages and T reg cells) and anti-tumorigenic populations (e.g., CD4⁺ T cells, CD8⁺ T cells, and DC cells) ^52^. Immunophenotyping analysis revealed a significant increase in cDC1 within USP18^KO^ tumours. cDC1s play a critical role in cross-presenting antigens and activating CD8⁺ T cells ^53–55^, suggesting enhanced priming of cytotoxic T lymphocytes. Consistently, we observed a significant increase in CD8⁺ T cells in USP18-deficient tumours, indicating a strengthened cytotoxic response against tumour cells. In addition to cytotoxic T cells, other key immune populations associated with antitumor immunity were also elevated. T helper cells (Th1) and pro-inflammatory macrophages, both central to orchestrating pro-inflammatory and immune-activating responses ^56–58^, were significantly increased in USP18^KO^ tumours. The elevation of Th1 cells and pro-inflammatory macrophage populations further highlight the coordinated activation of innate and adaptive immunity following USP18 depletion in tumours. Moreover, additional immune cell subsets, including neutrophils, B cells, antigen-presenting cells (monocyte-derived dendritic cells; moDCs), and NK cells, were also elevated in USP18^KO^ tumours. Together, these findings highlight USP18 as a negative regulator of anti-tumour immunity and demonstrate that loss of USP18 function in cancer cells enhances the immunogenicity of the tumour microenvironment in a TNBC model

USP18 functions as deISGylase, as well as a negative regulator of the IFN receptor complex ^26,28,33^. Several studies have used point mutations to generate enzymatically dead USP18 inhibition, including C61A (mouse), C64S (human), and C64R/C65R (human) ^28,30,32,34,37,59^. In addition, mutation of I60N selectively impairs the scaffolding function, without affecting the deISGylase activity. This was shown to be sufficient to enhance the IFN response in cancers ^28^. In this study, we use C64R/C65R (human) and C61R/C62R (mouse) as models for USP18 inhibition. Interestingly, the mutation phenocopies the loss of USP18, suggesting that the mutation inhibits the enzymatic and non-enzymatic function of USP18. Our data suggests that the introduction of larger R residues distorts a loop close to the active site, which may be required for the binding to STAT2 or IFNAR2. The IFN response was increased in USP18^C64R/C65R^ despite the augmented expression of mutant USP18, suggesting the complete abrogation of the negative feedback function. These findings demonstrate that targeting the catalytic site is a viable strategy to inhibit both the enzymatic and non-enzymatic functions of USP18. It is conceivable that an active site binder identified by high-throughput screening as an inhibitor of USP18 enzymatic activity could induce structural alterations similar to the C64R/C65R mutation and inhibit the scaffolding function as well. Screening for USP18 inhibitors using a scalable ISG15-Rhodamine-cleavage assay ^28,33^ is a more straightforward approach than screening for a USP18-STAT2 disruptor and able to identify bifunctional molecules. Together, these findings demonstrate that USP18 inhibition can enhance anti-tumour immune responses and promote a more immunologically active (“hot”) tumour microenvironment. This provides a strong rationale for developing USP18 inhibitors as novel immunomodulatory agents in cancer therapy.

## Acknowledgements

The authors gratefully acknowledge Walter Kolch for MDA-MB-231 cells and Simon Wilkinson for 4T1 cells for this study. Furthermore, we sincerely appreciate the Mass Spectrometry Facility and Flow Cytometry Facility at the Institute of Genetics and Cancer, University of Edinburgh, for their support.

## Authorship contribution statement

**Chartinun Chutoe**; Data curation, Formal analysis, Investigation, Methodology, Validation, Software, Visualization, Writing – original draft, review and editing. **Emily Webb**; Data curation, Formal analysis, Investigation, Methodology, Validation, Writing – original draft, review and editing. **Morwenna Muir**; Investigation and Methodology. **Fraser Laing**; Investigation and Methodology. **Valerie G Brunton**; Investigation, Methodology, Validation, Writing – original draft, review and editing. **Alexander von Kriegsheim**; Data curation, Formal analysis, Investigation, Methodology, Validation, Writing – original draft, review and editing.

