## Supplementary material for "USP18 Inhibition Enhances Type I Interferon Signalling and Immune Activation in the Tumour Microenvironment of Triple-Negative Breast Cancer": Supplemetary figures

* To whom correspondence should be addressed:

Alexander von Kriegsheim

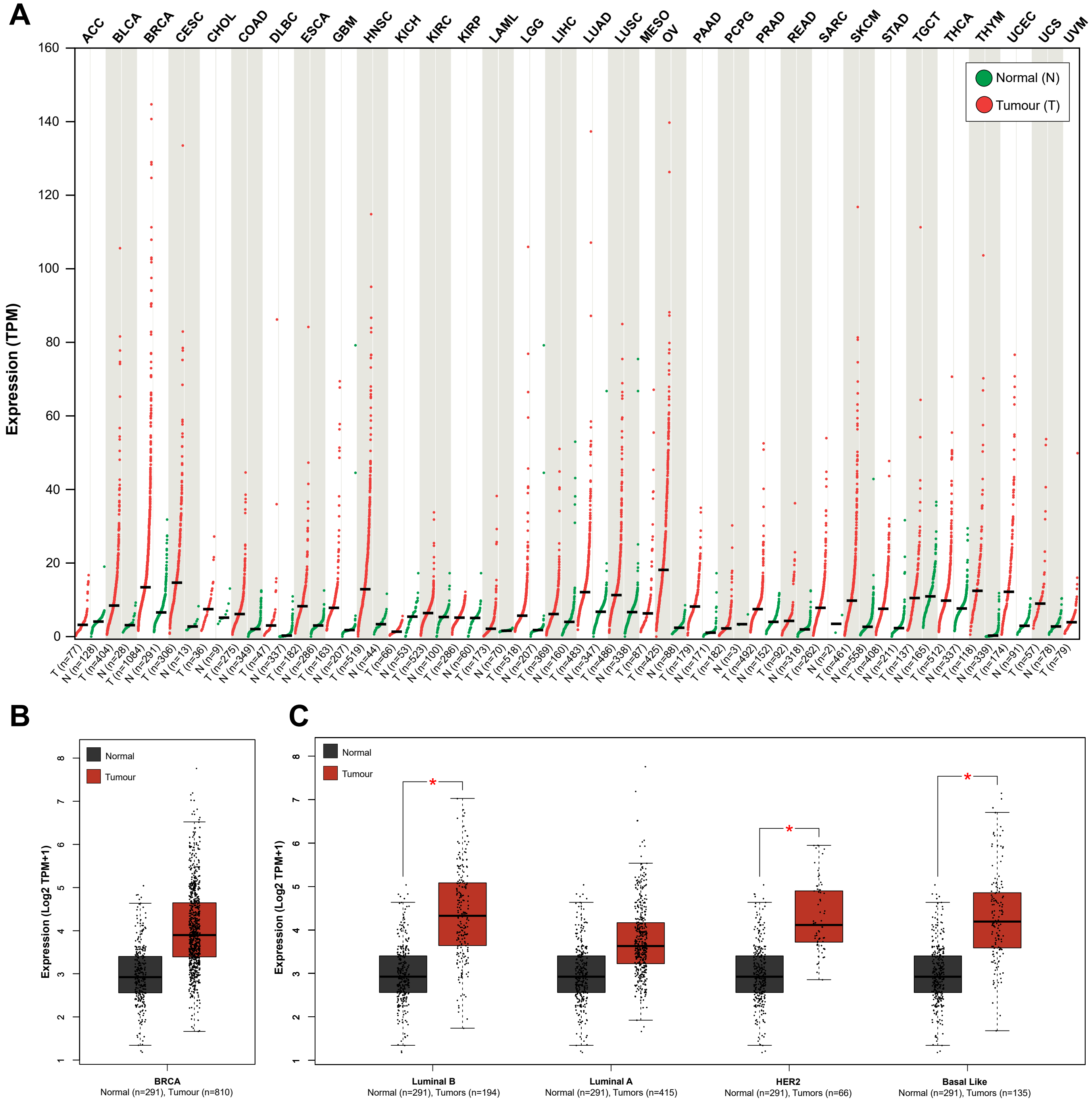


**Figure S1. USP18 was upregulated in breast cancer (BRCA).** Meta-analysis of RNA sequencing by using GEPIA2 showed **(A)** USP18 expression across different cancer types compared to normal tissues. **(B)** USP18 expression profile of breast cancer (BRCA) versus normal tissues (from fig. S1A), which further identify into **(C)** USP18 expression profile of each breast cancer subtypes. *p<0.05

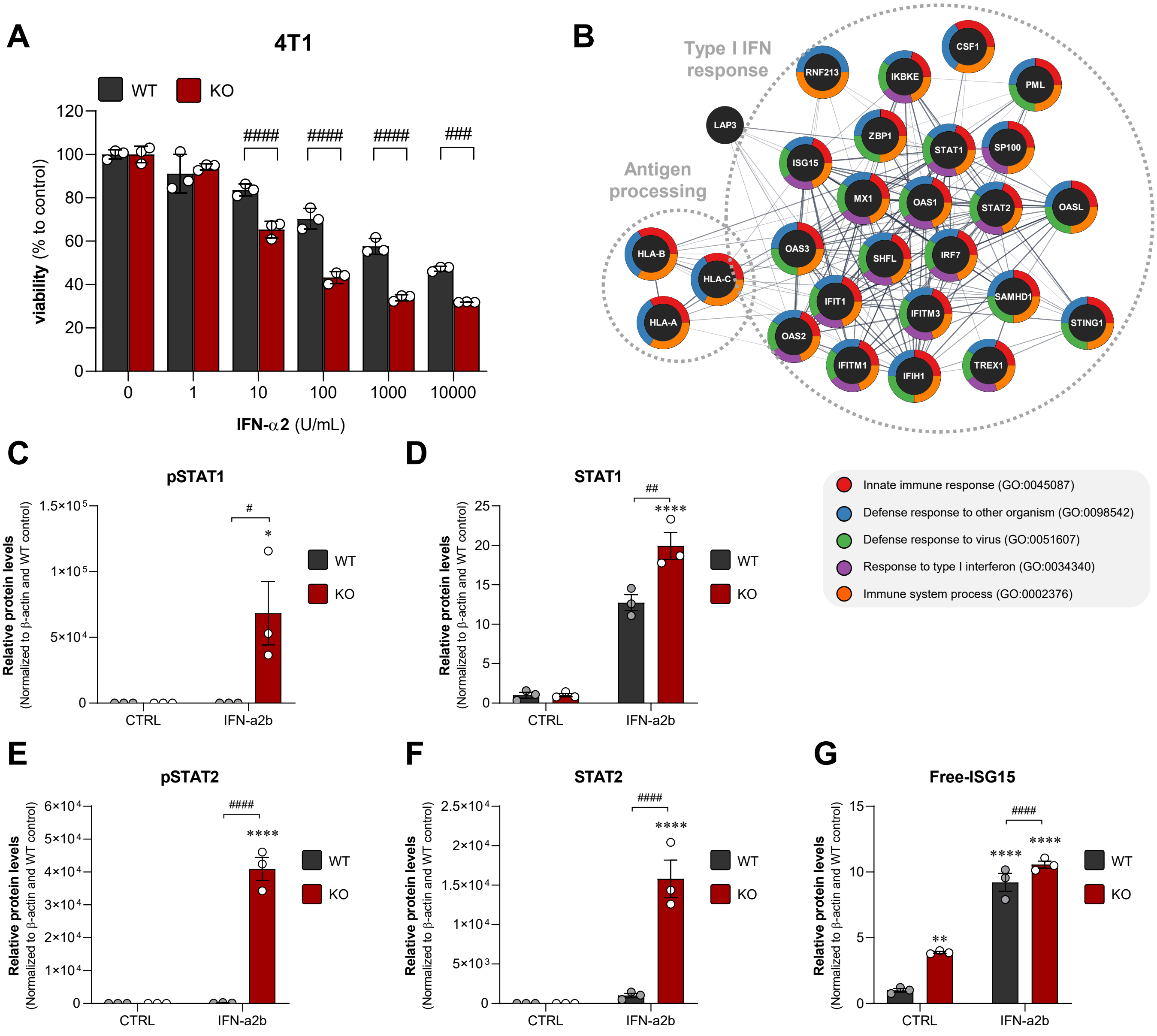


**Figure S2. USP18 depletion enhanced IFN response in breast cancer cells. (A)** Mouse triple-negative breast cancer (4T1) cell viability comparing USP18$^{WT}$ and USP18$^{KO}$ cells after 48 hours IFN-α2 exposure. **(B)** The ISG-protein signature was revealed by proteomic analysis. The quantification of ISG-protein signatures by immunoblot (shown in Fig.1); **(C)** p-STAT1, **(D)** STAT1, **(E)** p-STAT2, **(F)** STAT2, and **(G)** free-ISG15. Graphs showed mean of 3 independent biological replicates ± SEM. *$p<0.05$, **$p<0.01$, and ****$p<0.0001$ versus control, while #$p<0.05$, ##$p<0.01$, ###$p<0.001$, and ####$p<0.0001$.

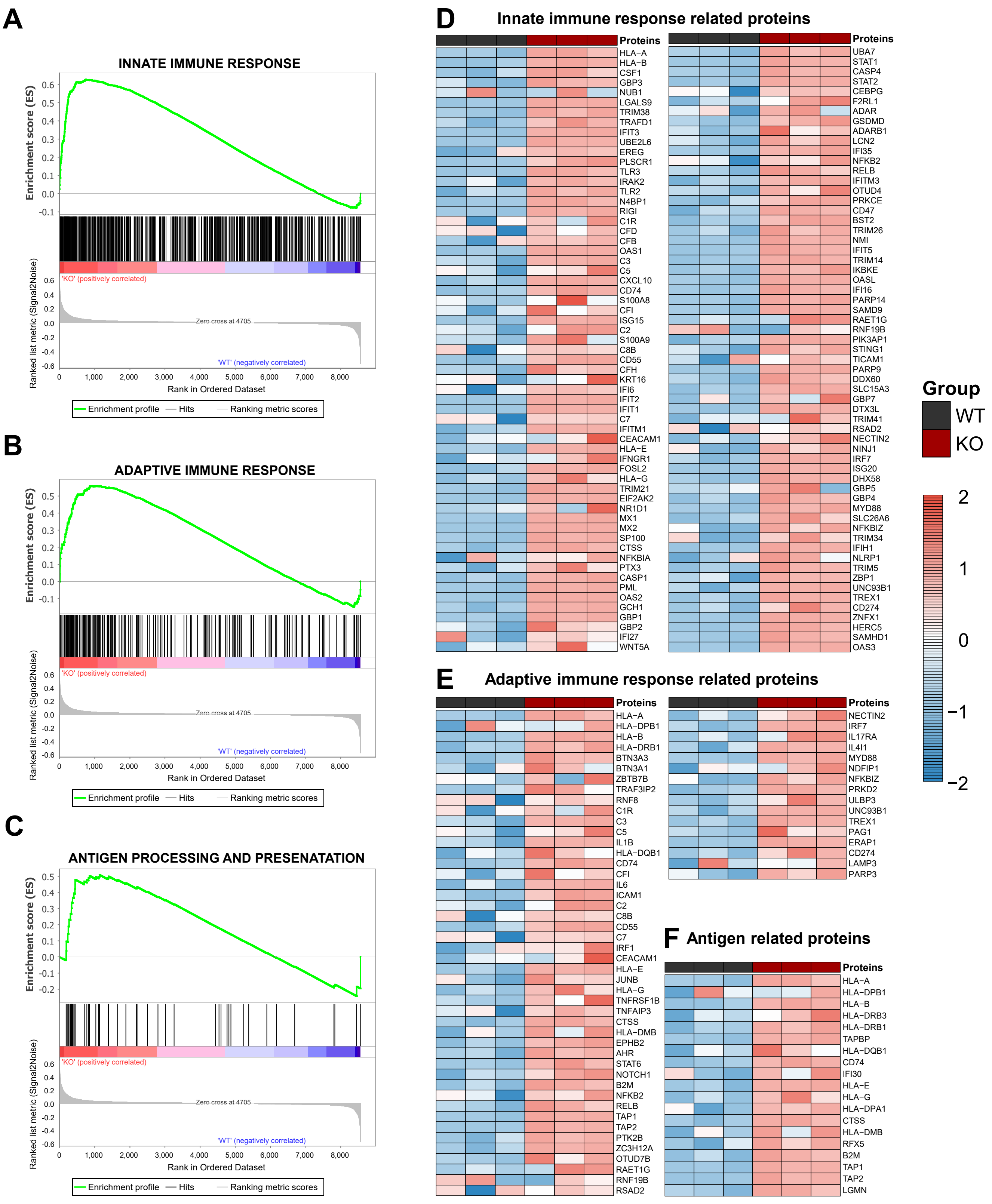


**Figure S3. Loss of USP18 enhanced immune response and cancer immunogenicity.** GSEA analysis of proteomic data illustrates **(A)** innate immune response, **(B)** adaptive immune response and **(C)** antigen processing and presentation. Heat map showed the significantly enriched proteins from GSEA analysis; **(D)** innate immune response **(E)** adaptive immune response, and **(F)** antigen processing and presentation. Proteomic data was acquired by 3 independent biological replicates.

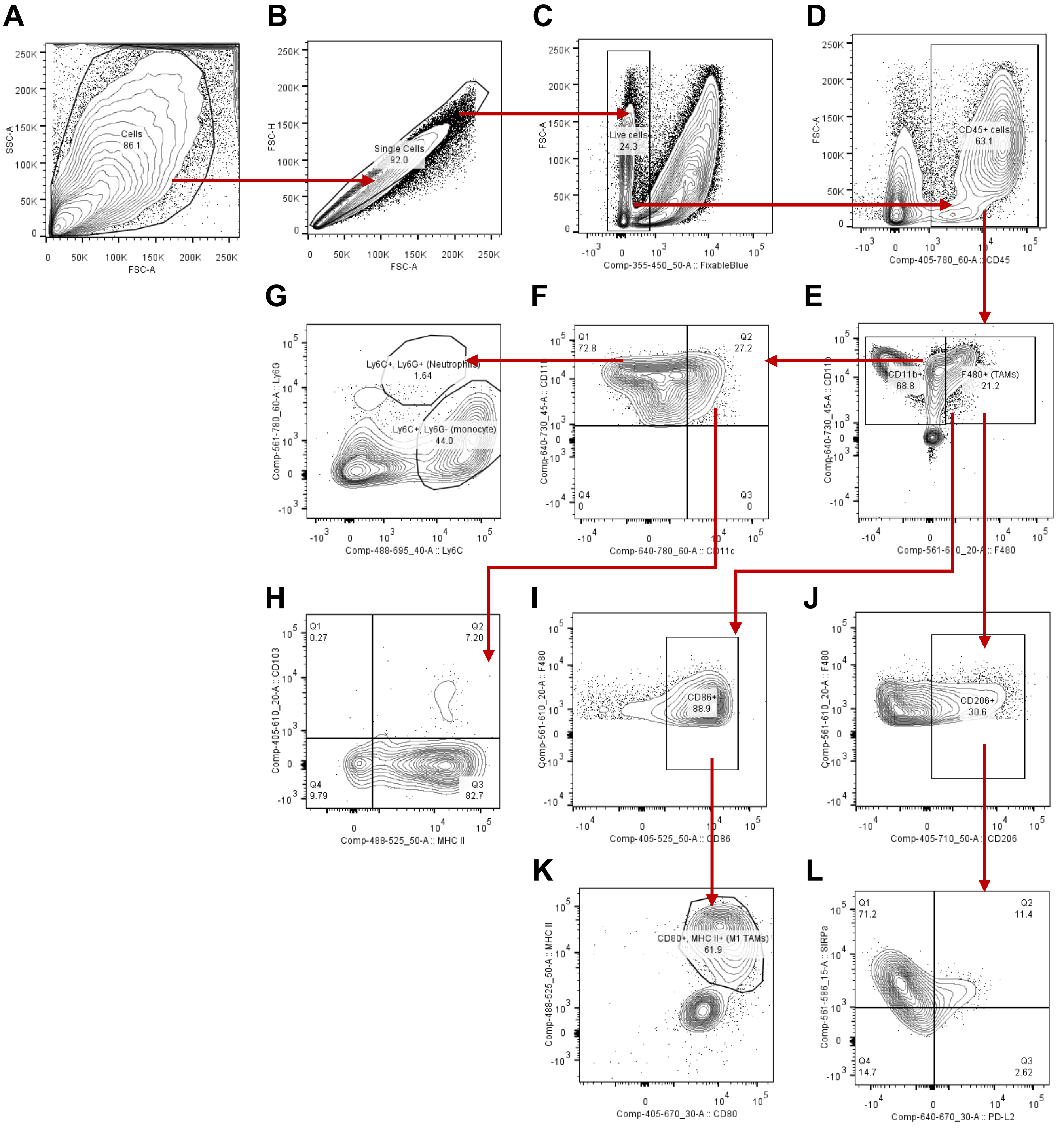


**Figure S4. Gating hierarchy for myeloid cells analysis from 4T1 tumours.** **(A)** Total cell event. **(B)** Identifying single .cells. **(C)** Identifying living cells. **(D)** Gating for immune ($CD45^+$) cells. **(E)** Gating for CD11b and F4/80 subpopulation. **(F)** Gating for CD11b and CD11c. **(G)** Gating for Ly6C and Ly6G subpopulation. **(H)** Gating for CD103 and MHC-II subpopulation. **(I)** Identifying $CD86^+$, $F4/80^+$. **(J)** Identifying $CD206^+$, $F4/80^+$. **(K)** Gating for $CD80^+$ (M1). **(L)** Gating for M2. The analysis was done further for other markers.

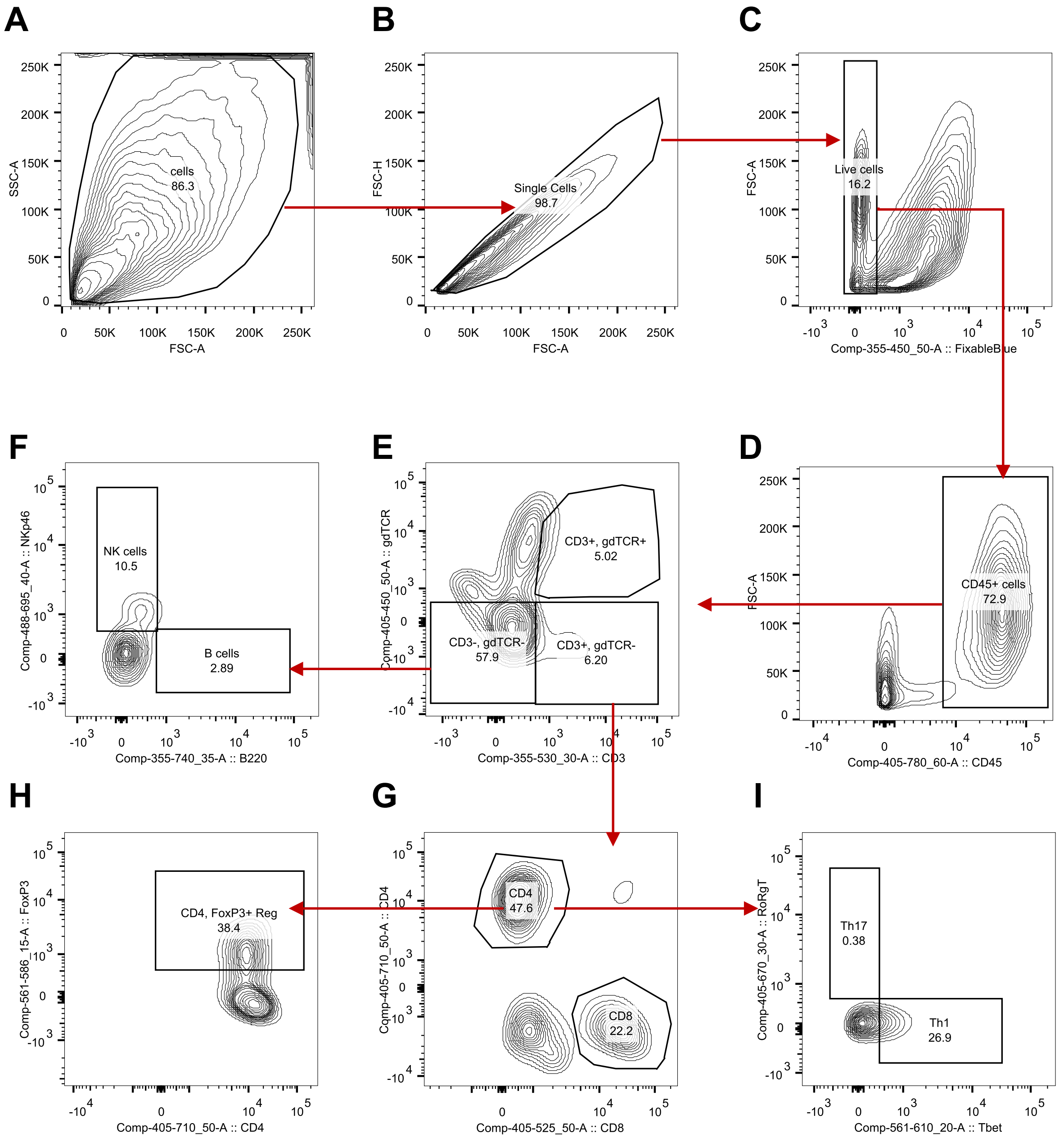


**Figure S5. Gating hierarchy for T cells analysis from 4T1 tumours. (A)** Total cell event. **(B)** Identifying single .cells. **(C)** Identifying living cells. **(D)** Gating for immune (CD45$^+$) cells. **(E)** Gating for CD3 and γδ TCR subpopulation. **(F)** Gating for NK cells (NKp46$^+$) and B cells (B220$^+$). **(G)** Gating for CD4 T cells (CD4$^+$) and CD8 T cells (CD8$^+$). **(H)** Gating for FoxP3$^+$. **(I)** Gating for Tbet$^+$ (Th1 cells) The analysis was done further for other marker as well as intracellular marker such as FoxP3 and Granzyme B etc.

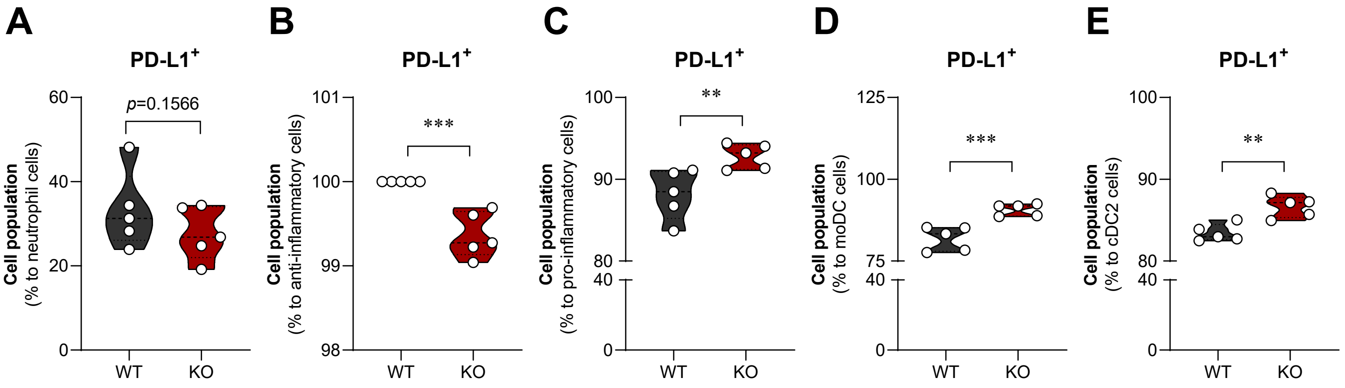


**Figure S6. Differential PD-L1 expression across myeloid subtypes in the tumour microenvironment.** Graphs showed a proportion of PD-L1$^{+}$ cells across myeloid subtypes including **(A)** neutrophils, **(B)** anti-inflammatory macrophage, **(C)** pro-inflammatory macrophage, **(D)** moDCs, and **(E)** cDC2. **p<0.01 and ***p<0.001 versus WT tumours.

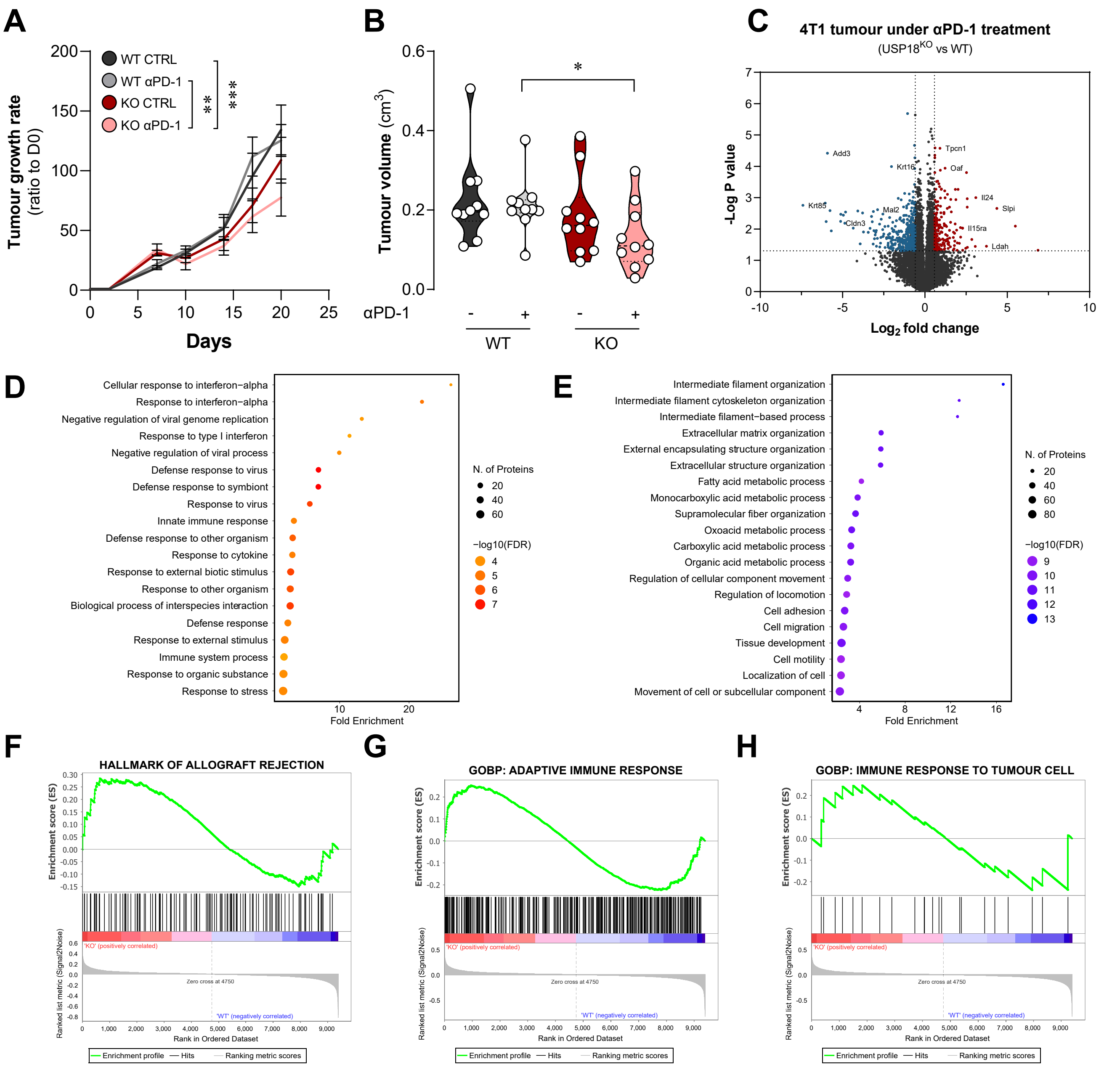


**Figure S7. αPD-1 treatment and USP18 depletion tumours *in vivo*. (A)** Tumour growth rate. **(B)** Tumour volume at day 20 after implantation. Proteomic study of tumour showed an alteration as **(C)** volcano plot, which changed **(D)** upregulated pathways and **(E)** downregulated pathways. Gene set enrichment analysis showed positive correlation of USPKO tumour under αPD-1 treatment to **(F)** hallmark of allograft rejection, **(G)** adaptive immune response, and **(H)** immune response to tumour cell. Graphs showed mean ± SEM. **p<0.01, and ****p<0.0001.

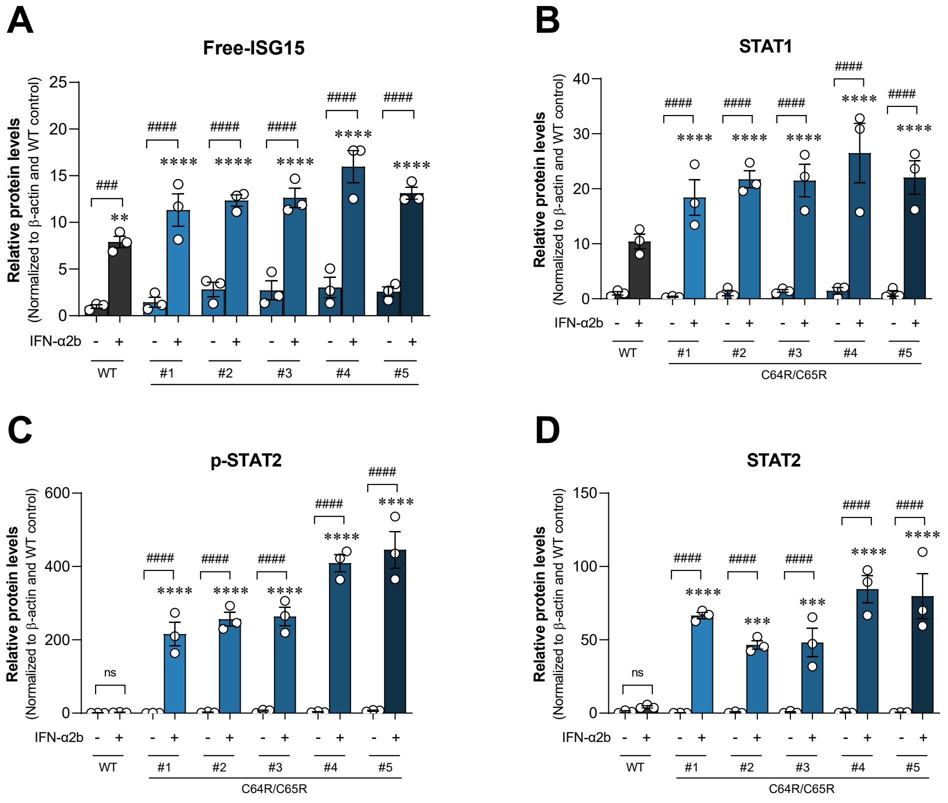


**Figure S8. Immunoblot analysis of IFN-signature proteins (shown in Fig. 5).** The quantification of band intensity of **(A)** free-ISG15, **(B)** STAT1, **(C)** p-STAT2, **(D)** STAT2. **(E)** Graphs showed mean of 3 independent biological replicates ± SEM. (ns=no statistically difference), **p<0.01, ***p<0.001, and ****p<0.0001 versus WT control, while ###p<0.001, and ####p<0.0001.

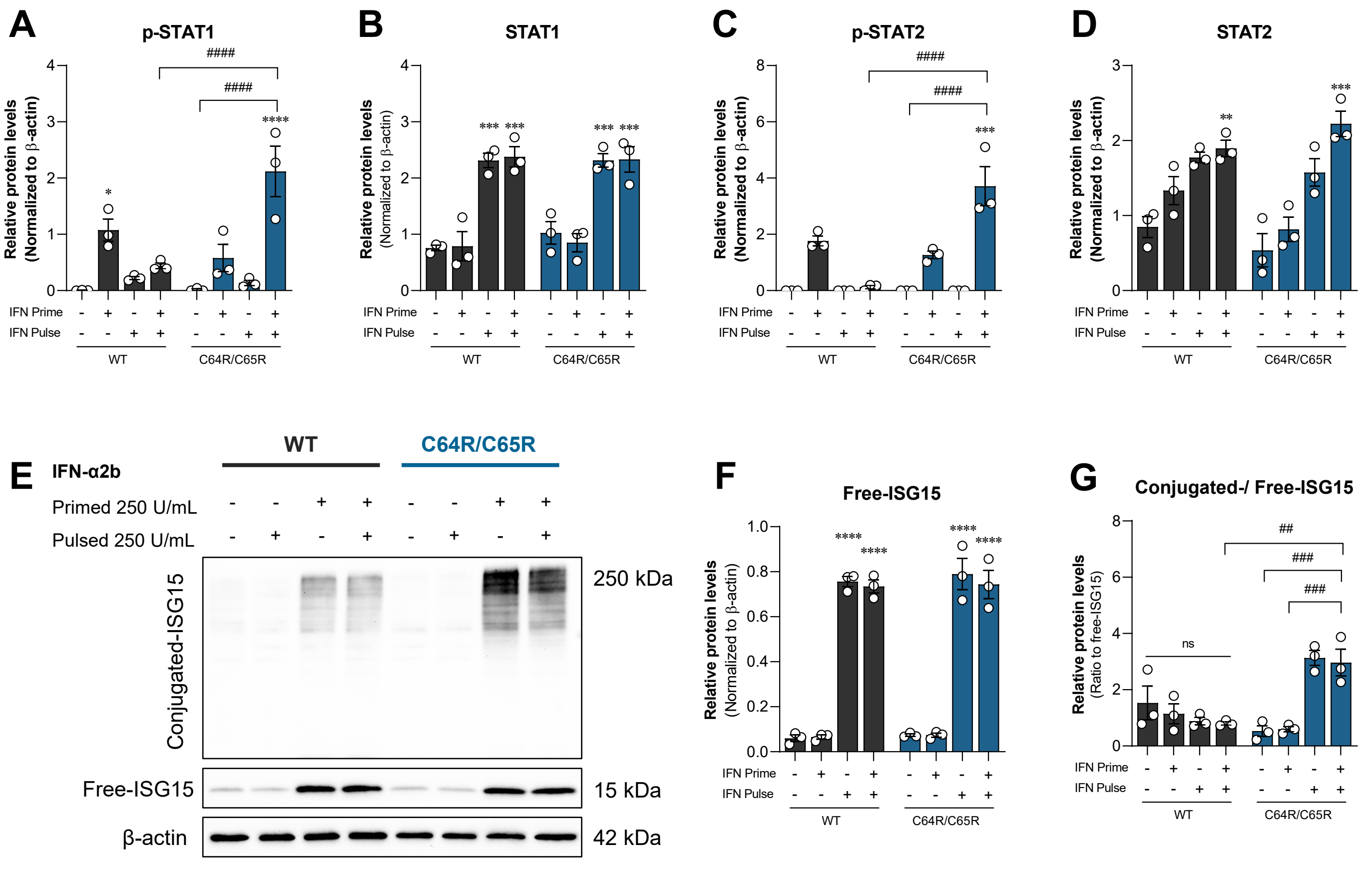


**Figure S9. Immunoblot analysis of IFN-signature proteins prime/pulse experiment (shown in Fig. 6).** The quantification of band intensity of **(A)** p-STAT1, **(B)** STAT1, **(C)** p-STAT2, **(D)** STAT2. **(E)** The represented blot of ISG15 and the quantification of **(F)** free-ISG15 and **(G)** conjugated-/free-ISG15 ratio. Graphs showed mean of 3 independent biological replicates ± SEM. *p<0.05, **p<0.01, ***p<0.001, and ****p<0.0001 versus control.

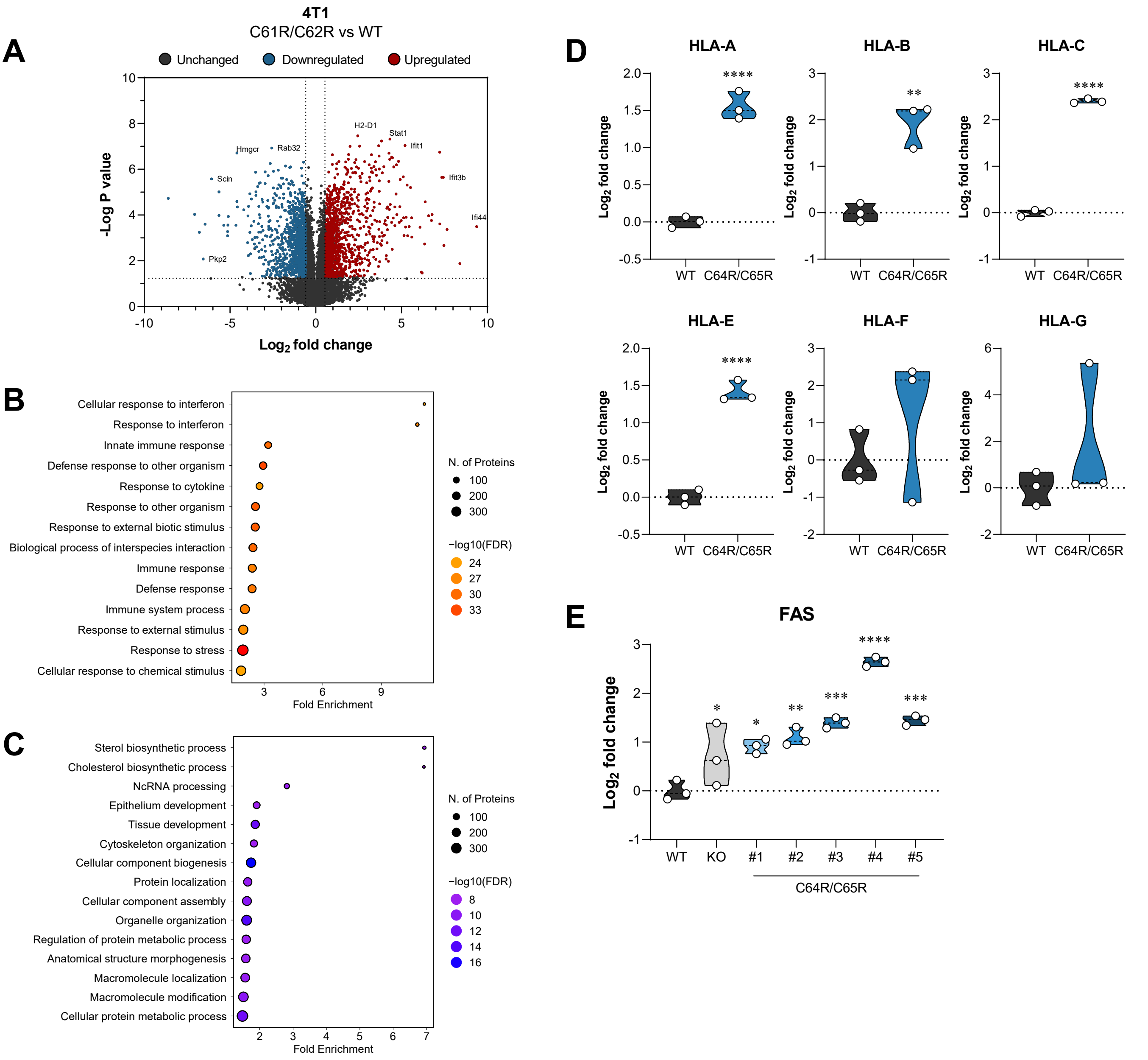


**Figure S10. Proteomic studies of catalytically mutated USP18.** The quantification of band intensity of **(A)** Volcano plot of USP18$^{C61R/C62R}$ 4T1 cells. 4T1 proteome alteration showed enrichment plots of **(B)** upregulated protein and **(C)** downregulated proteins. **(D)** MHC-I expression in USP18$^{C64R/C65R}$ MDA-MB-231 cells. **(E)** FAS receptor expression in USP18$^{C64R/C65R}$ MDA-MB-231 cells. Graphs showed mean of 3 independent biological replicates ± SEM. (ns=no statistically difference), p>0.05. *p<0.05, **p<0.01, ***p<0.001, and ****p<0.0001 versus control, while ##p<0.01, ###p<0.001, and ####p<0.0001.

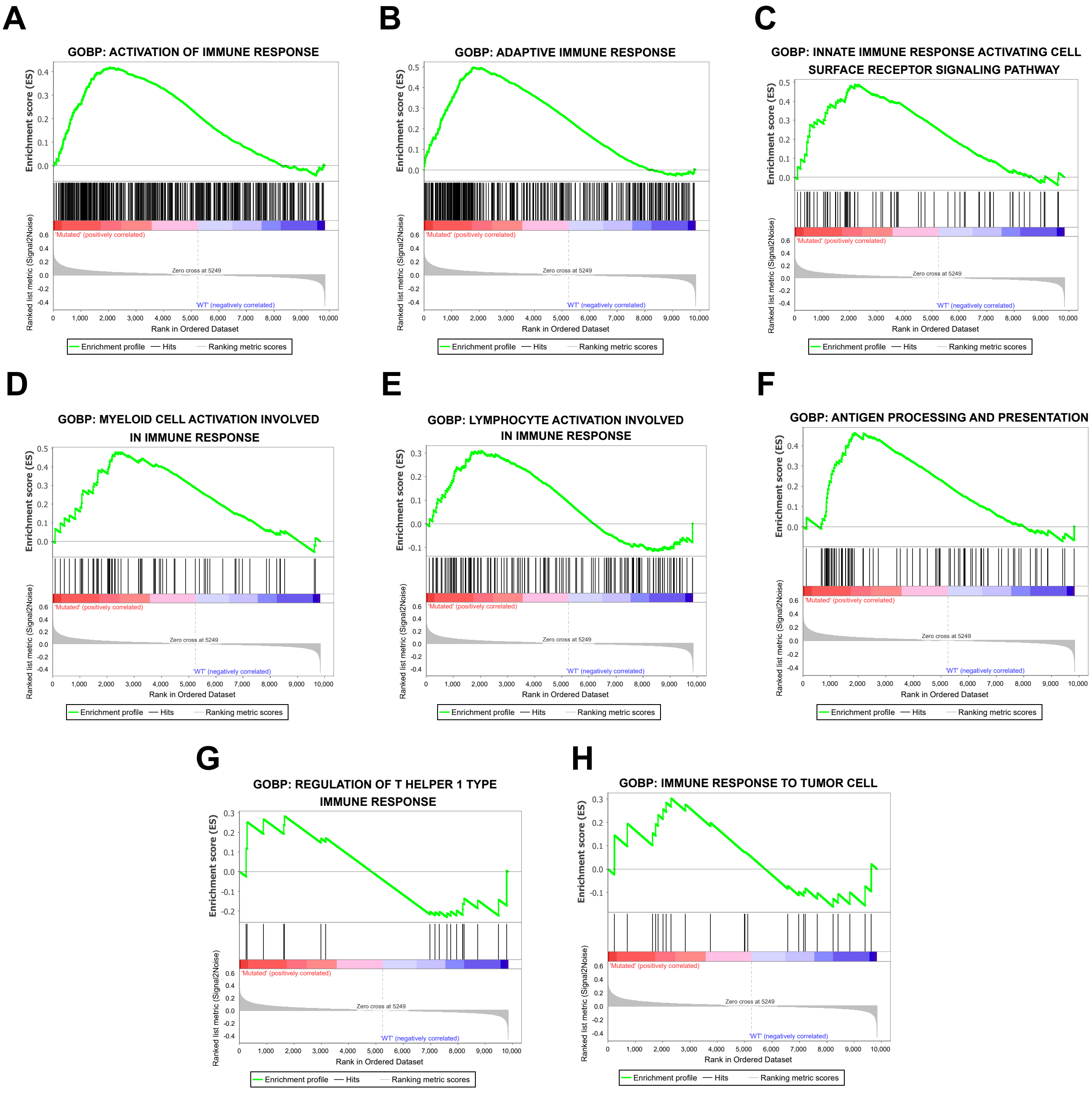


**Figure S11. Gene set enrichment analysis showed that C61R/C62R tumours positively correlated with immune activation related pathways. (A)** Activation of immune. **(B)** Adaptive immune response. **(C)** Innate immune response activating cell surface receptor signalling. **(D)** Myeloid cell activation involved in immune response. **(E)** Lymphocyte activation involved in immune response. **(F)** Antigen processing and presentation. **(G)** **(H)** Immune response to tumour cells.
