## Supplementary material for "USP18 Inhibition Enhances Type I Interferon Signalling and Immune Activation in the Tumour Microenvironment of Triple-Negative Breast Cancer": Supplemetary Table

* To whom correspondence should be addressed:

Alexander von Kriegsheim

**Table S1.** The Table shows crRNAs and HDR templates used in this study.

| ***USP18 knock out studies*** | | |
|---|---|---|
| **Targets** | **Design ID** | **Sequence** |
| Human | Hs.Cas9.USP18.1.AA | CTTCACCCGGATCGTATACA |
| Human | Hs.Cas9.USP18.1.AD | CTTTTCTTCAAGATCTGCCG |
| Mouse | Mm.Cas9.USP18.1.AA | ACGGCACGTTGTATTTCTGC |
| Mouse | Mm.Cas9.USP18.1.AK | AGGCACTGAACGAGCTCCGT |
| ***USP18 knock in studies*** | | |
| **Targets** | **Design ID** | **Sequence** |
| Human | Hs.Cas9.USP18.1.AB | CATTACGAACACCTGAATCA |
| Mouse | Mm.HC9.CCHP3899.AA | ACACTTGAAGCAAGGAGTTA |
| ***HDR templates*** | | |
| Human | TCCCCTTATAGGCCTGGTTGGTTTACACAACATTGGACAGACCAGGAGGCTTAACTCTCTGATTCAGGTGTTCGTAATGAATGTGGACTTCACCAGGATATT | |
| Mouse | TTCAATATCATTCTGAAGTCCATGTTCATCATGAACACTTGAAGCAAGGAGTTAAGCCTCCTCGTCTGTCCGATGTTGTGTAAACCAACCAGACCTACAACATGAAGAGGGAAAAAAA | |

**Table S2.** The table shows list of antibodies used for immune cell profiling from tumour and lymph nodes.

| Cell types | Markers | Fluorophores | Suppliers | Catalog number |
|---|---|---|---|---|
| **Leukocytes** | NKp46 | PerCP-Cy5.5 | BioLegend | 137610 |
| | CD4 | BV711 | BioLegend | 100549 |
| | CD8 | Brilliant Violet 510™ | BioLegend | 100751 |
| | FoxP3 | PE | BioLegend | 126404 |
| | γδ TCR | Brilliant Violet 421™ | BioLegend | 118119 |
| | Tbet | PE-Dazzle 594 | BioLegend | 644828 |
| | Granzyme B | FITC | BioLegend | 372206 |
| | CD45 | BV786 | BD Biosciences | 564225 |
| | PD-1 | BV605 | BioLegend | 135219 |
| | TIM-3 | APC-Fire750 | BioLegend | 119738 |
| | CD3 | BUV 469 | BD Biosciences | 569671 |
| | B220 | BUV 737 | BD Biosciences | 612838 |
| **Myeloid** | CD45 | BV786 | BD Biosciences | 564225 |
| | CD11b | APC-R700 | BD Biosciences | 564985 |
| | F4/80 | PE/Dazzle™ 594 | BioLegend | 123146 |
| | Ly6C | PerCP/Cy5.5 | BioLegend | 128012 |
| | Ly6G | PE-Cy7 | BioLegend | 127618 |
| | CD86 | Brilliant Violet 510™ | BioLegend | 105039 |
| | CD206 | Brilliant Violet 711™ | BioLegend | 141727 |
| | CD80 | BrillViolet 650 | BioLegend | 104731 |
| | PD-L2 | APC | BioLegend | 107210 |
| | CD103 | Brilliant Violet 605™ | BioLegend | 121433 |
| | SIRPa | PE | BioLegend | 144012 |
| | MHC II | FITC | BioLegend | 107606 |
| | PD-L1 | BUV 737 | BD Biosciences | 568826 |
| | CD11c | APC-Fire750 | BioLegend | 117352 |
